# Affordable 96-parallel SPPS: hardware and software add-on for automated liquid handling platforms

**DOI:** 10.64898/2026.11.09.765483

**Authors:** Grigory L. Kozhemyakin, Ivan K. Chudinov, Mikhail S. Nosov, Nikita M. Baraboshkin, Angelina A. Lebedeva, Oleg V. Fedorov

## Abstract

ParSynt-P is a hardware and software add-on that converts a liquid-handling robot already available in a proteomics laboratory (Tecan Freedom EVO) into a parallel plate-format peptide synthesizer for generating libraries of candidate peptide standards for targeted blood plasma proteomics. The manifold is milled from polytetrafluoroethylene and has ports for an inert gas and a vacuum line; a standard commercially available 96-well filtration plate is secured to it with an airtight seal. The software layer generates robot worklists from a table of peptide sequences and controls switching of the N₂/vacuum lines, including pulsed gas mixing. The system performs up to 96 syntheses per run, and coupling of one residue takes 94 min. Workability of this 96-well format was verified on three fully loaded plates. The uniformity of synthesis conversion across the plate was measured quantitatively using a characterized NAT/SIL standard pair. The add-on cost $351, while the synthesis cost is $0.23 per coupling and $1.14 per milligram of crude product, an order of magnitude below the market price of custom synthesis.

## Introduction

### Context and need

Quantitative mass spectrometry of blood plasma proteins analyzes peptides rather than proteins: a target protein is measured from the signal of its proteotypic tryptic peptides, while synthetic peptides of the same sequence provide the metrological basis of the measurement as calibrants for external calibration curves and as isotope-labeled internal standards. This requires not only the peptides included in the panel itself, but also a considerably larger pool of candidates: the suitability of a particular sequence as a standard is established empirically by screening synthesized peptide libraries [1–3]. Thus, a typical targeted plasma proteomics panel covering dozens of proteins requires tens to hundreds of individual peptides [4, 5], in small amounts, from micrograms to milligrams per sequence, sufficient for constructing calibration curves and developing mass spectrometry (MS) methods. The need to synthesize a large number of peptides rapidly can be addressed in two ways: 1) ordering peptides commercially from a specialist organization, where the cost depends on the number and length of the peptides and may amount to tens of dollars per sequence, while delivery takes weeks, making iterative adjustment of the panel composition (replacing a peptide, adding an isoform, testing an alternative charged variant) slow and expensive; 2) in-house synthesis using a dedicated instrument, a peptide synthesizer. This eliminates dependence on a supplier, but the instrument itself costs tens of thousands of dollars [6, 7] and, in most configurations, permits one to a few syntheses in parallel, so the panel is assembled through successive runs.

At the same time, the critical component of a synthesis platform, precise, position-specific dispensing of small reagent volumes, is already part of the standard equipment of proteomics laboratories in the form of automated liquid handling stations. In this work, we show that purchasing a plate-format peptide synthesizer can be avoided: an existing robotic liquid handling platform needs only a specialized module and a software layer to provide the elements missing for solid-phase peptide synthesis (SPPS): an airtight reaction vessel with an inert atmosphere in the 96-well format, drainage of liquids through a filter bottom, and mixing.

### Automation of SPPS

Solid-phase peptide synthesis (SPPS), introduced by R. B. Merrifield [8], transformed peptide assembly into a cyclic process of ‘deprotection → washing → coupling → washing’ that is inherently amenable to automation; modern Fmoc chemistry and its reagent toolkit are systematized in methodological reviews [9, 10] that establish a de facto standard protocol, the same protocol implemented in this work.

Automation is developing along three distinct lines: accelerating an individual cycle, expanding the chemical capabilities of a single run, and parallelizing multiple syntheses.

The time required for a single coupling has been reduced through flow synthesis, from rapid flow-based SPPS [11] and a fully automated setup with a cycle of approximately a few minutes per residue [12] to the assembly of full-length protein chains [13]. The same principle is also established in commercial instruments with one to a few reactors: microwave protocols have progressed to complete elimination of washing within the cycle [14], while induction heating at 25–90 °C has been implemented on the PurePep Chorus [15]. Expansion of chemical capabilities proceeds independently of sequence synthesis speed: the Chemputer platform describes the entire protocol in the formal language χDL, from resin swelling to peptide precipitation, and incorporates post-synthetic modifications (ring-closing metathesis, copper-catalyzed azide–alkyne cycloaddition, native chemical ligation) into the same continuous cycle [16]. All these systems optimize either the time or the chemical diversity of a single chain; the number of parallel syntheses is limited to a few, and dozens of different sequences are obtained through several successive runs.

A second practical solution is to increase the number of parallel couplings, and two approaches have historically developed in this area.

The first is SPOT synthesis and peptide arrays, in which hundreds to thousands of short sequences are assembled at individually addressable positions on a common support [17]; modern implementations achieve densities above 17 000 spots per cm² through laser-based transfer of monomers in a solid matrix [18]. This level of multiplexing is unattainable in reactor formats, but the material remains on the support and is usually not cleaved into solution. Cleavable SPOT formats have come closest to the task of producing individual peptides: Glyco-SPOT yields parallel glycopeptide libraries in microgram quantities suitable both for solution assays and for microarray printing [19]. The limitation of this technology lies less in quantity than in the support and its use: specialized equipment, a membrane or spot rather than a standard well, and a separate excision and cleavage procedure.

The second approach comprises instruments that perform full SPPS in parallel in many separate reactors, and it is this category to which the comparable systems discussed below belong. Modern commercial instruments of this class already operate in a plate format: MultiPep 2 (CEM/Intavis) accommodates four 96-well reactors and yields 384 peptides per run, Syro I and Syro II (Biotage) offer 4 × 96 and 6 × 96 positions, and Vantage and Apex 396 (Aapptec) offer 96 and 4 × 96 positions [20]. Until recently, the denser 384-well format had been implemented only using 4 mm membrane disks in a MultiPep RS holder, yielding approximately 30 nmol of peptide per well, enough for screening protein–protein interactions but not for preparative purposes; adapting a 384-well plate to conventional polystyrene resin required modification of the instrument and yielded 1536 peptides per synthesis at a mean scale of 0.5 μmol per well and crude purity above 80 % [20]. The limitation of this approach is not multiplexing but price and closed architecture: the cost of commercial valve-based and robotic synthesizers is estimated at more than $30 000 [21] and more than $50 000 [6], which prompted the development of open-source solutions: the Arduino-and Python-based PepSy, with an assembly cost below $4000 [21], and an extremely low-cost ($300-600) version based on consumer peristaltic pumps [22].

The approach conceptually closest to the present work is based on integrating a solid-phase peptide synthesis module into the basic configuration of a standard laboratory liquid handling station. The TaPSy system was built as an add-on module for a Tecan platform and performs parallel synthesis of up to 12 peptides, dividing the tasks between the pipetting robot (dispensing) and a separate N₂/vacuum valve circuit (purging, mixing, liquid removal) [6]. This architecture eliminates the main cost item, because the dispensing device is already available, but in the published work the system is limited to 12 positions using individual cartridges. This defines the niche of the present work: not record-breaking multiplexing, since plate formats have already been implemented in dedicated instruments, but the full 96-well format without purchasing a synthesizer, namely a compact add-on for a liquid-handling robot already installed in the laboratory, with a standard removable filtration plate and an open software layer for protocol generation.

### Synthetic peptides in targeted proteomics

The practical significance of parallel synthesis of individual peptides is directly linked to the needs of quantitative proteomic analysis. The human blood plasma proteome has an exceptionally broad dynamic concentration range and remains a central focus of biomarker discovery [23, 24]. Reliable quantification of specific proteins uses targeted mass spectrometric methods, selected/multiple reaction monitoring (SRM/MRM) and its high-resolution counterpart, parallel reaction monitoring (PRM), which rely on measuring the signals of proteotypic peptides (unique to a protein and reproducibly detectable) [25, 26]. Selection of such peptides has been formalized computationally [27], and a large-scale resource of validated targeted assays covering the entire human proteome is compiled in the Human SRMAtlas [28].

A key component of these methods is synthetic peptides that reproduce proteotypic sequences of the target proteins and are used as standards, including isotope-labeled forms for absolute quantification (the AQUA strategy) [29, 30]. Targeted blood plasma proteomics panels include tens to hundreds of proteins and therefore require the same, and sometimes a greater, number of peptides; their production is a typical task for parallel plate-format SPPS. The applied part of the present work addresses precisely this task.

## Materials and methods

### Design and fabrication of the manifold, holder, and reagent reservoirs

The manifold, holder, and reagent reservoirs were designed using Autodesk Fusion computer-aided design software (Autodesk Inc., USA). The manifold and reagent reservoirs were machined from a solid polytetrafluoroethylene (PTFE-F4) blank using a computer numerical control milling machine. The holder was fabricated from polypropylene filament by fused deposition modeling on a Flashforge Adventurer 5M 3D printer (Jinhua Flashforge Technology Co., Ltd., China). The materials were selected for their chemical resistance to piperidine bases, DMF, and trifluoroacetic acid.

### SPPS chemistry (synthesis cycle protocol)

The protocol was tested on a commercial PurePep Chorus automated peptide synthesizer (Gyros Protein Technologies, USA), for which the chemistry used is standard. Synthesis followed a solid-phase peptide synthesis protocol using amino acid derivatives protected at the α-amino group with a 9-fluorenylmethoxycarbonyl (Fmoc) group. Wang resin preloaded with N-α-Fmoc-N-ε-Boc-L-lysine (Fmoc-Lys(Boc)-Wang) (Sigma-Aldrich), with a loading capacity of 0.35 mmol/g, was used as the solid phase; 25 mg was loaded per reactor, corresponding to 8.75 μmol.

Each synthesis cycle comprised three Fmoc deprotection treatments with a 20 % solution of 4-methylpiperidine (Sigma-Aldrich) in N,N-dimethylformamide (DMF) (Supelko), followed by washing of the resin with DMF, coupling of the next Fmoc-protected amino acid residue using 2-(7-aza-1H-benzotriazol-1-yl)-1,1,3,3-tetramethyluronium hexafluorophosphate (HATU) (Sigma-Aldrich) and N,N-diisopropylethylamine (DIPEA) (Sigma-Aldric) for 30 min, and two subsequent DMF washes. Reagent solutions were dispensed in 250 μL portions and washes in 2250 μL portions; the solution concentrations were 210 mM for the Fmoc-amino acid, 200 mM for HATU, and 420 mM for DIPEA, corresponding to 52.5, 50, and 105 μmol, or 6.0, 5.7, and 12.0 equivalents relative to the resin loading capacity (molar ratio Fmoc-amino acid : HATU : DIPEA = 1 : 0.95 : 2.00); the activating reagent was used in a slight deficit relative to the Fmoc-amino acid.

The established protocol was transferred, without changes to the chemistry or solution concentrations, to a Freedom EVO 150 automated liquid handler (Tecan, Switzerland) equipped with the “ParSynt-P” add-on. Synthesis was performed in a 96-Well Filtration Plate, 2.0 mL/well, 5 μm frit/well (Biocomma). The resin loading was 10 mg of Fmoc-Lys(Boc)-Wang per well (3.5 μmol). Dispensing volumes were 100 μL for each reagent solution, 180 μL for the deprotection step, and 900 μL for washing. The deprotection and coupling cycles were repeated sequentially according to the specified amino acid sequence.

Analytical standards were synthesized by SPPS on a PurePep Chorus automated peptide synthesizer (Gyros Protein Technologies, USA) using amino acid derivatives with Fmoc-protected α-amino groups. Two forms of the same sequence were synthesized: the natural form (NAT) on Fmoc-Lys(Boc)-Wang resin (0.35 mmol/g, 17.5 μmol) and the isotope-labeled form (SIL) on Fmoc-Lys(Boc)-[¹³C₆,¹⁵N₂]-TCP resin (Iris Biotech) (0.21 mmol/g, 10.5 μmol). Fmoc groups were removed from the first amino acid residue on the support and from the growing peptide chain using a 20 % solution of 4-methylpiperidine in DMF. Fmoc-amino acid coupling (6 equiv.) was performed in the presence of DIPEA (12 equiv.) using the activating system 2-(1H-benzotriazol-1-yl)-1,1,3,3-tetramethyluronium tetrafluoroborate / 1-hydroxybenzotriazole (TBTU/HOBt) (Sigma-Aldrich) (6 equiv./3 equiv.); the coupling reaction lasted 30 min. After coupling, unreacted amino groups were capped with propionic anhydride (10 equiv.) in the presence of DIPEA (10 equiv.) in DMF for 10 min. The resin was washed with DMF between successive synthesis steps.

Peptide cleavage from the solid phase with simultaneous side-chain deprotection was performed in a mixture of trifluoroacetic acid (TFA) (Sigma-Aldrich) : 3,6-dioxa-1,8-octanedithiol (DODT) (Sigma-Aldrich) : triisopropylsilane (TIS) (Sigma-Aldrich) : anisole (Sigma-Aldrich) : water (Milli-Q) at a volume ratio of 183:5:2:5:5 for 3 h at RT; 1 mL of the mixture was used per 10 mg of resin. After incubation, the solution was collected and ice-cold diethyl ether (Sigma-Aldrich) was added. The resulting suspension was held for 30 min at −80 °C and centrifuged for 10 min at 7000g. The pellet was dried under vacuum and dissolved in 1 mL of 5 % acetonitrile in water containing 0.01 % TFA.

After cleavage and analysis, both forms were purified by preparative reversed-phase HPLC on an ÄKTA pure 25 system (Cytiva, USA) using an Agilent ZORBAX SB-C18 column, 9.4 × 250 mm, 5 μm (Agilent, USA), with a linear gradient from 5 to 40 % mobile phase B over 20 min; the mobile phases were A, 0.01 % TFA in water, and B, 0.01 % TFA in acetonitrile. The collected fractions were lyophilized and dissolved in 1.0 mL of 5 % acetonitrile in water containing 0.01 % TFA.

### Analytical methods

Crude products were analyzed by reversed-phase HPLC with simultaneous spectrophotometric and mass spectrometric detection. An LC-40 liquid chromatograph (Shimadzu, Japan) with a UV–visible spectrophotometric detector coupled to a Triple Quad 3500 triple-quadrupole mass spectrometer (SCIEX, USA) was used. Separation was performed on a Luna C18 column, 4.6 × 250 mm, 5 μm (Phenomenex, USA), at 30 °C. The mobile phases were A, 0.01 % trifluoroacetic acid in water, and B, 0.01 % trifluoroacetic acid in acetonitrile; a linear gradient from 5 to 100 % B over 20 min was applied at a flow rate of 0.9 mL/min; the injection volume was 1 μL, and spectrophotometric detection was performed at 220 nm. Mass spectra were acquired in first-quadrupole scanning mode (Q1 scan) over the m/z range 250-2000 in positive-ion mode.

The completeness of synthesis was assessed as the fraction of the target peptide peak area in the total chromatographic peak area at 220 nm, taken as the purity of the crude product. The identity of the target peak was confirmed from the mass spectrum acquired in Q1 scan mode, using the m/z values of the singly and doubly charged peptide ions.

Mass spectrometry-grade acetonitrile (Concord, China), mass spectrometry-grade trifluoroacetic acid (Sigma-Aldrich, USA), and deionized water produced using a Milli-Q purification system (Merck, Germany) were used to prepare mobile phases and dissolve samples.

### Amino acid analysis of the standards

Peptide standard concentrations were determined by amino acid analysis with liquid chromatography–mass spectrometry detection, following the method of Qasrawi et al. [31]. The method is based on acid hydrolysis of the peptide to free amino acids followed by quantification by HPLC–MS/MS in selected reaction monitoring mode; calculations use an external calibration curve with signal normalization to an isotope-labeled internal standard.

All steps (preparation of calibration solutions, addition of peptide samples and internal standard, acid hydrolysis, solvent removal, and dissolution of hydrolysates) and HPLC–MS/MS analysis were performed directly in polypropylene PCR strips.

Amino Acids Mix Solution (Supelco, USA) was used as the external standard. A commercial mixture of isotope-labeled amino acids, Stable Isotope Labeled Amino Acid Mix Solution 1 (Supelco, USA), served as the internal standard. A nine-point calibration series was prepared over the range 20–2000 pmol/μL (20, 50, 100, 200, 500, 750, 1000, 1500, and 2000 pmol/μL) by adding calculated volumes of standard solutions at concentrations of 0.025, 0.25, and 2.5 mM; 20 μL of internal standard solution was added to every sample, including calibration solutions and peptide samples. The peptide aliquot volume was 10 μL.

After all components had been added, the samples were dried in a Centrivap vacuum concentrator (Labconco, USA) at 60 °C for 2 h. Each dry residue received 75 μL of 6 M hydrochloric acid prepared by diluting concentrated (37 %) hydrochloric acid with Milli-Q water; the tubes were sealed tightly and incubated in a dry heating block at 110 °C for 24 h. After hydrolysis, the samples were cooled to room temperature and dried again in the vacuum concentrator at 60 °C for 2 h. The dry residues were dissolved in 50 μL of 0.01 % aqueous trifluoroacetic acid, treated in an ultrasonic bath, and briefly centrifuged; the strips were sealed with a silicone mat and placed in the chromatographic system autosampler.

Amino acid analysis was performed on a Luna C18 column, 4.6 × 250 mm, 5 μm (Phenomenex, USA), at 30 °C and a flow rate of 0.9 mL/min using the following gradient (mobile phase B): 1 % for 1 min, a linear increase to 50 % by 6 min, a hold at 50 % until 11 min, a return to 1 % by 11.01 min, and equilibration until 16.01 min. Detection was performed in selected reaction monitoring mode with a Turbo Spray ionization source at the following settings: curtain gas 40, collision gas 7, ion spray voltage 5500 V, temperature 650 °C, ion source gas 1, 65, and ion source gas 2, 65. The transitions, retention times, and potentials for all measured amino acids and their isotope-labeled analogs are provided in S1 Table.

An individual calibration curve was constructed for each amino acid from the external standard series using the ratio of the amino acid peak area to the peak area of the corresponding isotope-labeled internal standard; the curve was fitted by least squares with 1/x weighting. The amino acid concentration in the sample was calculated from this curve, accounting for the sample volume and dilution factors at all sample preparation stages. Peptide concentration was calculated from the amino acid concentrations, taking into account the number of corresponding residues in the sequence; when a peptide contained several amino acids suitable for quantification, the calculation was performed independently for each, and agreement between the resulting values was used as an internal analytical control. For the isotope-labeled standard, lysine was excluded from the calculation because the hydrolysate contains Lys^+8^ and the Lys/Lys^+8^ ratio no longer reflects the analyte content. The other amino acids are unaffected and are suitable for both forms. Each sample was analyzed in duplicate.

### Measurement of peptide concentrations

Peptide concentrations in the samples were determined by HPLC–MS/MS in selected reaction monitoring mode (SRM). The calibration curve was constructed using a synthetic standard characterized by amino acid analysis and signal normalization to an isotope-labeled internal standard of the same sequence.

A 10 μL aliquot was taken from the peptide solution (1 mL of 5 % acetonitrile containing 0.01 % trifluoroacetic acid), and 10 μL of isotope-labeled standard working solution (1000 pmol/μL) and 180 μL of the same solvent were added; the final sample volume was 200 μL, the original solution was diluted 20-fold, and the internal standard concentration in the sample was 50 pmol/μL.

Working solutions of the heavy and light standards at 1000 pmol/μL were prepared by dilution of the stock solutions (5949.05 and 5715.39 pmol/μL, respectively). Calibration solutions were prepared by adding calculated volumes of light standard working solutions at concentrations of 10, 100, and 1000 pmol/μL, 20 μL of heavy standard working solution, and solvent to a final volume of 400 μL, giving eight calibration points with nominal concentrations of 2.5, 5, 10, 25, 50, 100, 250, and 500 pmol/μL at an internal standard concentration of 50 pmol/μL, the same as in the analyzed samples; a blank sample was also analyzed.

Separation was performed on an XTerra RP18 column, 3.0 × 150 mm, 3.5 μm (Waters, USA), at 30 °C and a flow rate of 0.5 mL/min using the following gradient: 5 % phase B for 1 min, a linear increase to 100 % by 11 min, a hold until 16 min, a return to 5 % by 16.01 min, and equilibration until 21 min; the mobile phases were A, Milli-Q water containing 0.01 % trifluoroacetic acid, and B, acetonitrile containing 0.01 % trifluoroacetic acid. Detection used a Turbo Spray ionization source at the following settings: curtain gas 40, collision gas 7, ion spray voltage 5500 V, temperature 650 °C, ion source gas 1, 65, and ion source gas 2, 65. For each peptide, four transitions from the doubly charged precursor ion were monitored for the light and isotope-labeled forms; transition parameters are provided in S2 Table. Quantitative processing used the single most intense transition, b6; the other three transitions were used to confirm identity. The calibration curve was constructed using eight levels, with three replicates per level, by least squares with 1/x² weighting. The peptide concentration in each well was calculated accounting for the 20-fold sample dilution and the dissolution volume (1 mL), yielding the amount of peptide in the well and, consequently, the yield for each reaction position on the plate. Quantification was performed for one peptide in the model set, AVRPGYPK.

## Results and discussion

### A minimal add-on for a liquid-handling robot: design, synthesis protocol, and software layer

#### Reaction plate

A standard 96-Well Filtration Plate with 2 mL wells and frits with a pore size of 5 μm was selected as the reaction vessel. This choice was based on three factors: 1) the 5 μm pores hold DMF, preventing it from draining spontaneously, so the reaction mixture remains in the well throughout incubation and is drained only when vacuum is applied; 2) nitrogen delivered from below forms numerous small bubbles that lift and mix the solid-support beads; 3) the standard plate is compatible with a 96-deep well collection plate (96-DWP), is replaced between syntheses, and requires no washing.

#### Manifold, gas line, and vacuum line

To enable liquid removal and purging of the reaction plate, a polytetrafluoroethylene (PTFE-F4) manifold was designed to match the dimensions of a standard 96-well filtration plate (134 × 94 × 40 mm) (S1 Fig). The assembly sequence is shown in Fig 1.

**Fig 1.**
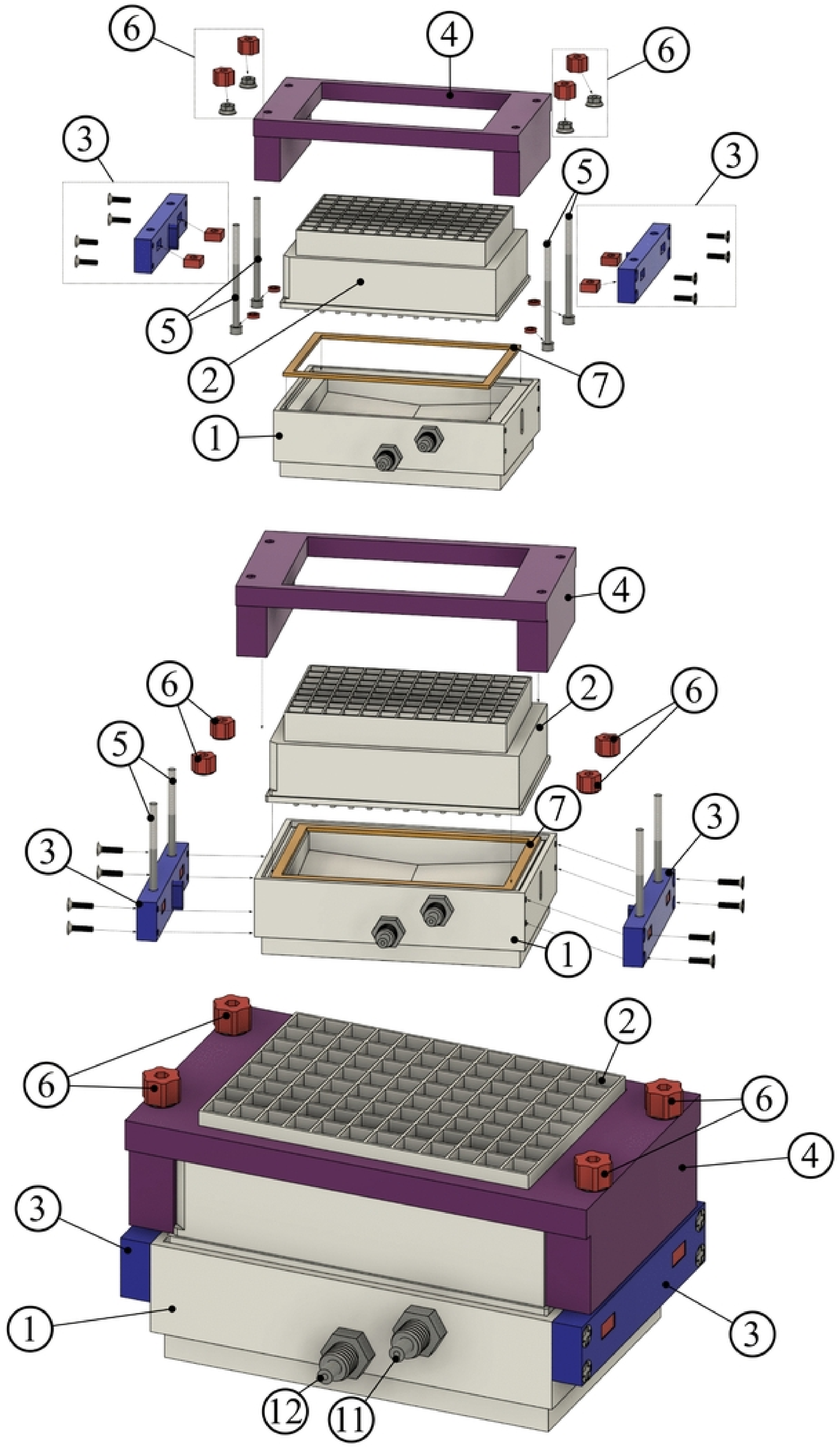
Assembly of the unit: manifold, filtration plate, and holder (items 1–7).

The unit is assembled as follows: the reaction plate (2) is placed on the manifold (1) over a sealing element (7), a 2 mm thick silicone gasket seated in a groove on the upper surface of the body, and held by an airtight clamping device (3): the holder (4) is tightened with wing nuts (6) on four threaded fasteners (5) (stainless steel M4x60 screws, d = 4.0 mm, l = 60 mm, overall length 64 mm; head h = 4.0 mm, d = 7.0 mm). The bottom of the internal cavity inclines toward the vacuum port, ensuring complete removal of drained reagents. The internal cavity of the manifold (8; Fig 2) communicates with two ports: the first (9) supplies inert gas, and the second (10) is used for vacuum removal. Fittings for the nitrogen supply line (11) and vacuum removal line (12) are screwed into the ports, with an internal tapered G1/4″ pipe thread and a hose barb for tubing with a 4 mm internal diameter; the vacuum fitting is located below the gas fitting and aligned with the lowest point of the sloping bottom. The tubing and fittings in both circuits are made of PTFE, which is chemically resistant to the solvents used.

**Fig 2.**
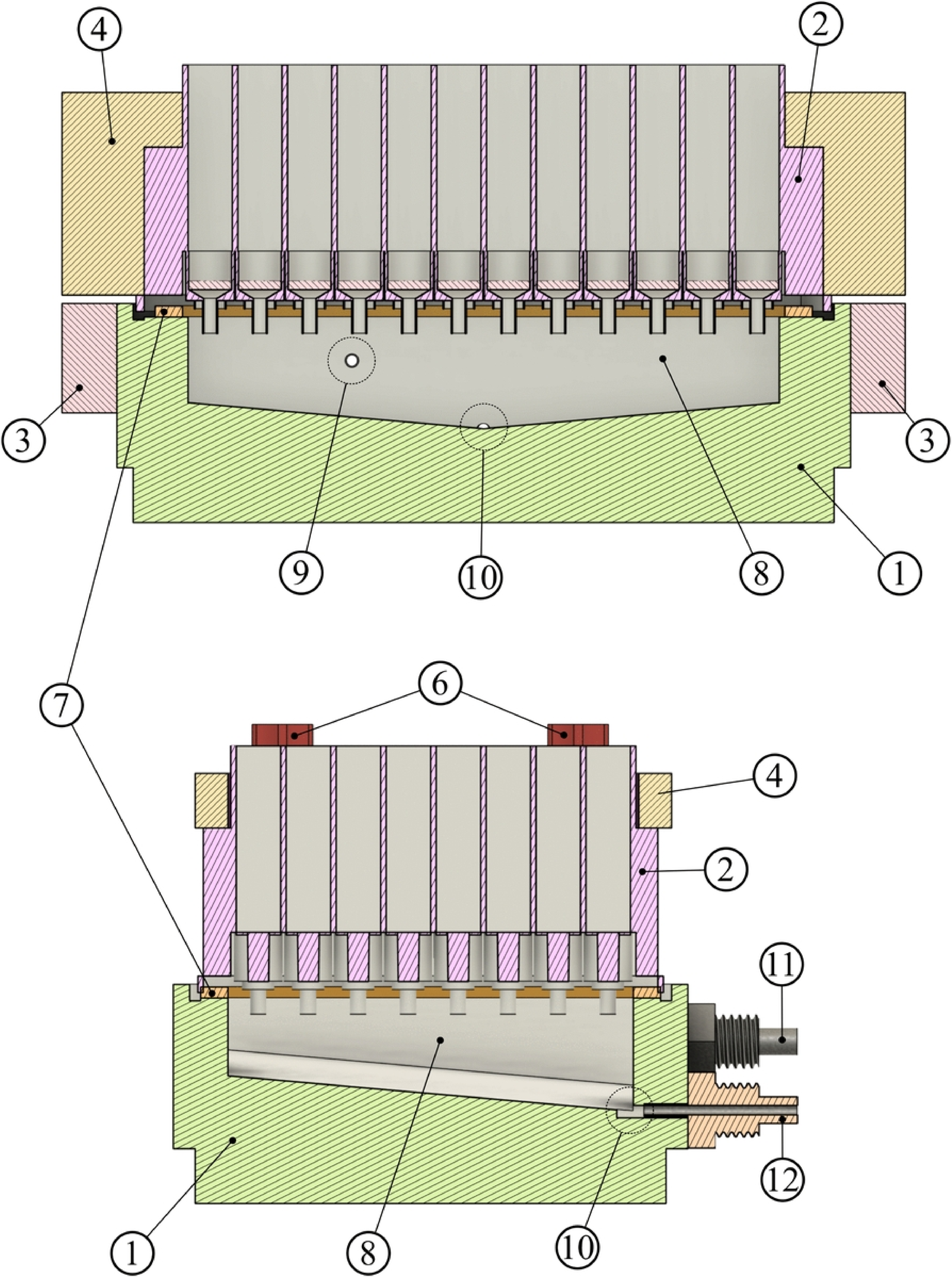
Manifold sections: internal cavity, ports, and fittings (items 8–12).

Draining is performed by a Wiggens C610 pump (China) through a 2500 mL collection vessel (19; Fig 3), a bottle with a standard GL 45 neck and holes drilled in its cap for fittings. Liquid is removed after each step for 30 s at the maximum vacuum attainable with the pump used.

**Fig 3.**
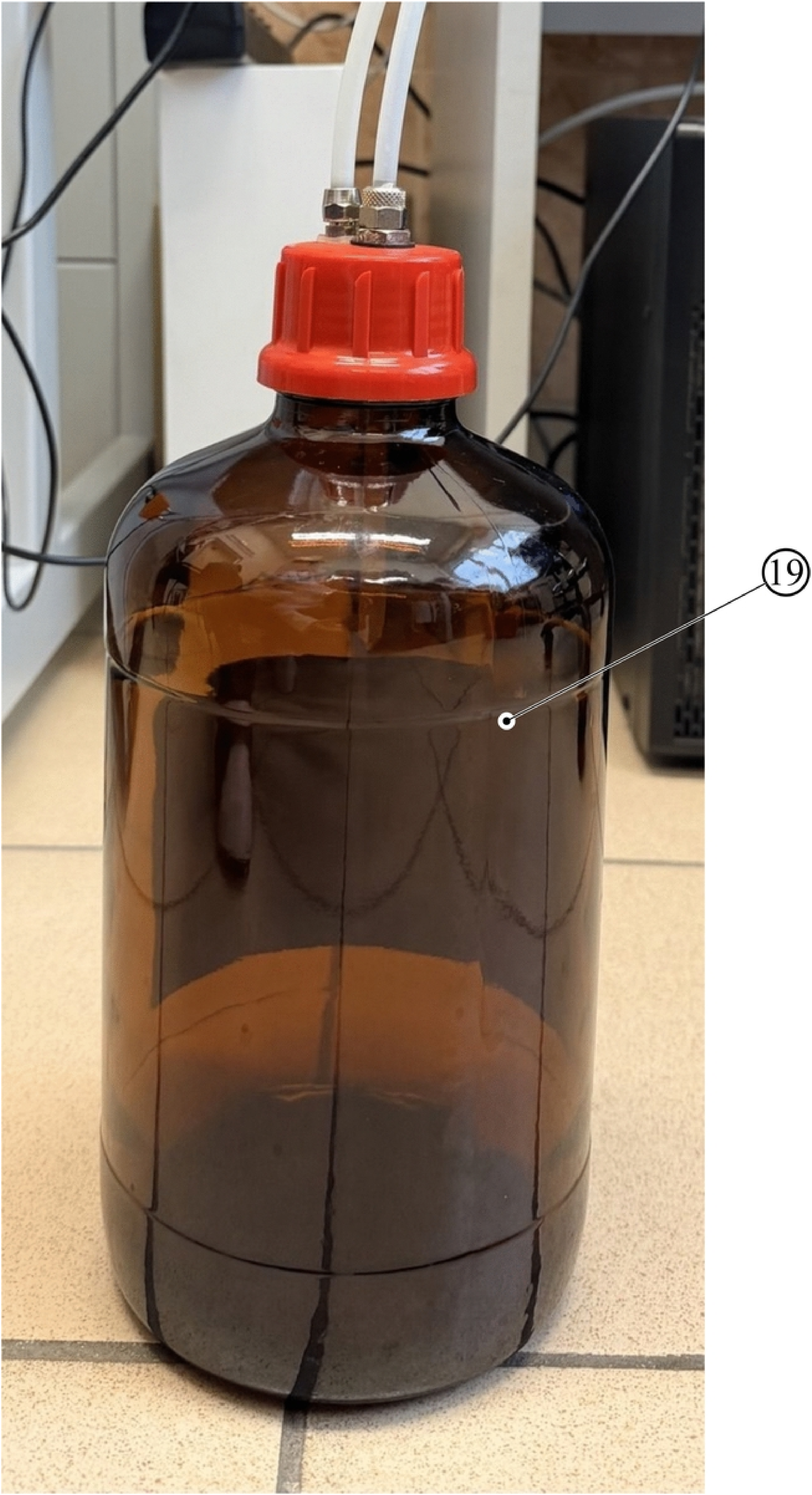
Collection vessel (item 19).

Gas supply and vacuum activation are controlled by two ELP08-040E4 solenoid valves (E-MC, China): the first (13) is installed in the nitrogen line and the second (14) in the vacuum line (Fig 4).

**Fig 4.**
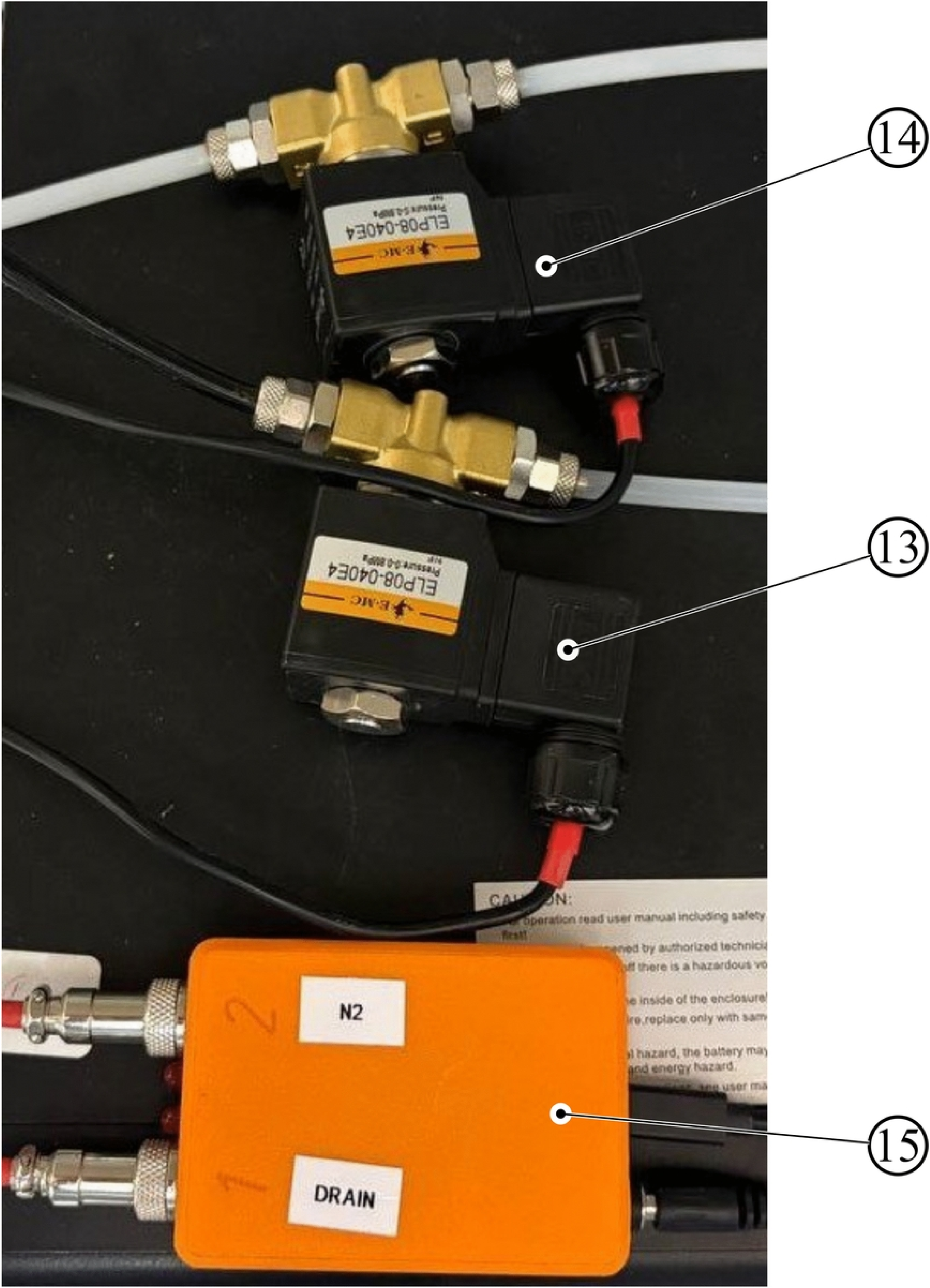
Valves and control unit (items 13–15).

Valve power is switched by a control unit (15) based on an STM32F103 microcontroller board (‘Blue Pill’) with two relay modules; the board is connected to the control computer via a USB cable (port rate 115 200 baud) and is recognized by the system as a virtual COM port (S1 File Cotroller_mod.zip). The circuit state is encoded by two bits: the high-order bit corresponds to the vacuum valve and the low-order bit to the gas valve: 00, both closed; 01, nitrogen line open; 10, vacuum line open; the combination 11 is not used in the synthesis protocol. Control uses Python scripts (control.py for line switching and stir.py for pulsed mixing by alternating commands 00 and 01); program execution is initiated by the liquid handling system, synchronizing valve switching with the dispensing protocol. The line pressure is set by the cylinder regulator (2 bar), and gas flow through the manifold is adjusted manually using a microvalve.

#### Minimal reagent set and deck layout

The reagent set is reduced to five items (Table 1), for which 200 mL PTFE reservoirs (16; Fig 5) were designed within the 96-DWP footprint to occupy standard deck slots: DMF for washing, deprotection solution (20 % 4-methylpiperidine in DMF), DIPEA, and HATU. Each reservoir is closed with a lid (17) containing a row of liquid aspiration holes (18) at a 9 mm pitch for a single pass of the 8-channel manipulator, so all eight tips enter the common volume simultaneously while the rest of the solution surface remains covered. Amino acids are placed in a standard 2 mL 96-DWP.

**Fig 5.**
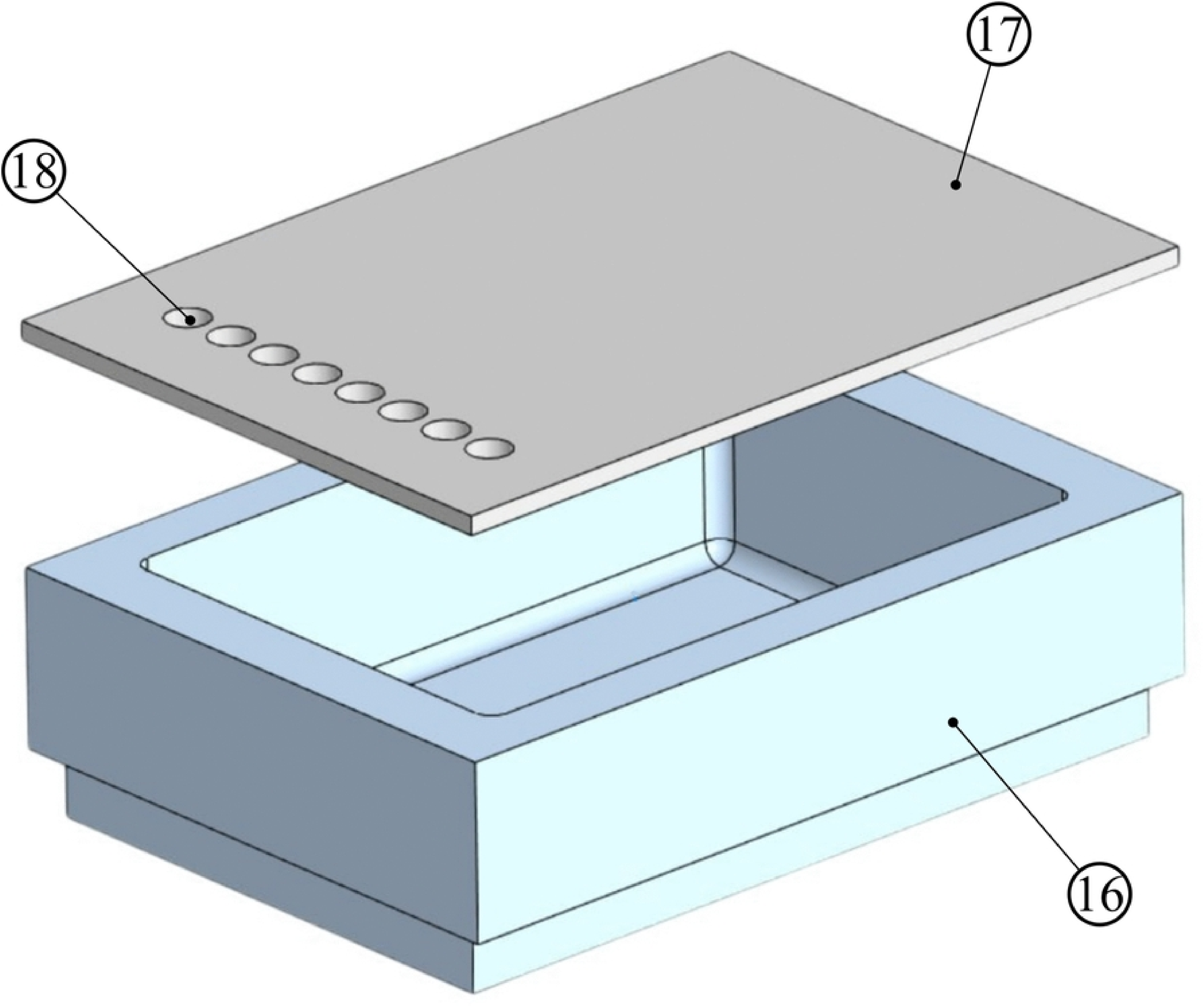
Reagent reservoir with lid (items 16–18).

The add-on installed in an automated liquid-handler system (20) is shown in Fig 6: the manifold with the clamped plate, reagent reservoirs, and valve unit occupy standard deck slots, and the gas line runs from the cylinder regulator on the left.

**Fig 6.**
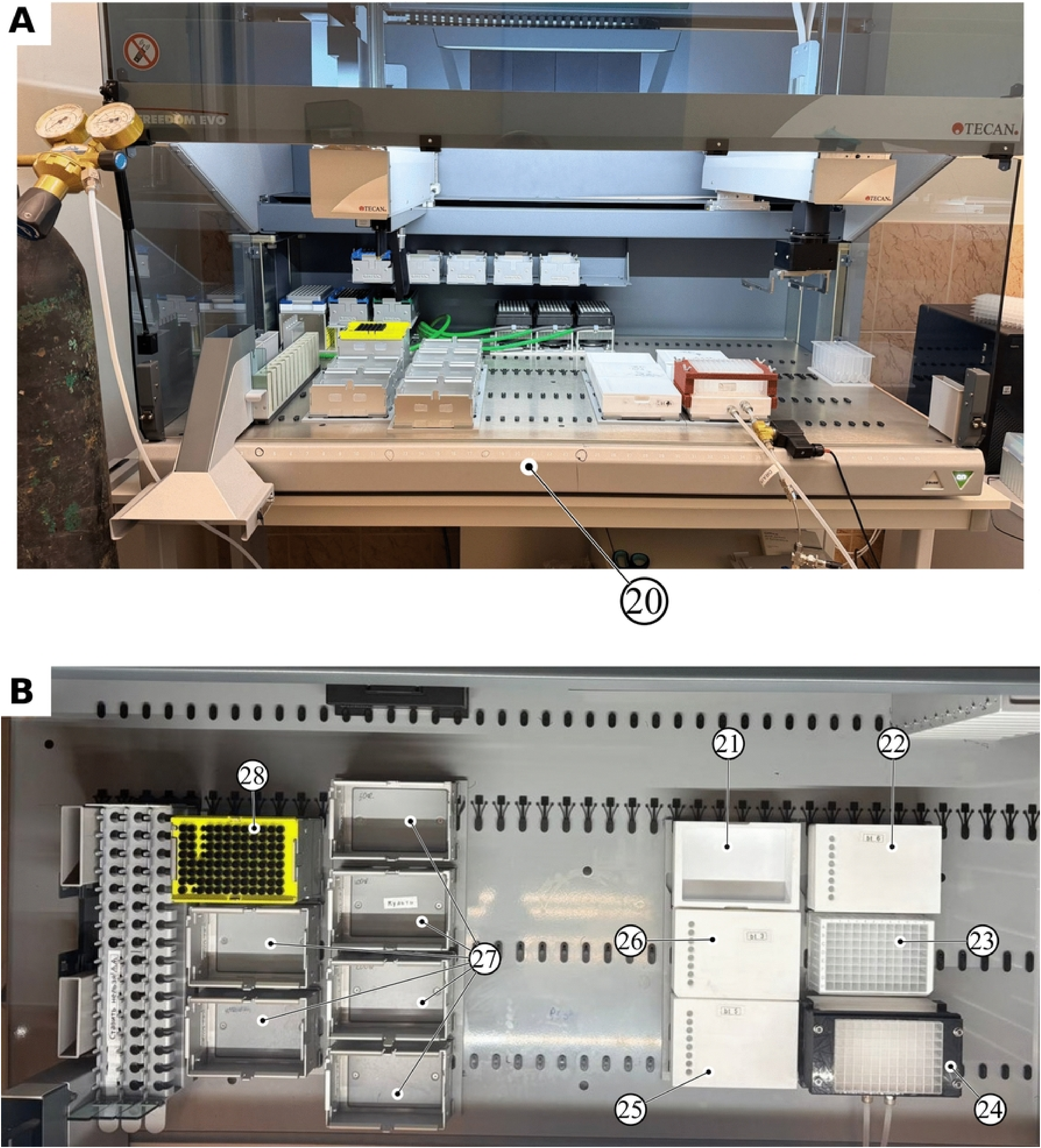
(A) The add-on installed in the liquid-handling robot (20) and (B) a top view of the working area. (21), DMF; (22), HATU; (23), Fmoc-amino acids; (24), reaction plate; (25), DIPEA; (26), 20% 4-methylpiperidine; (27), 200 μL tips; (28), 1000 μL tips.

**Table 1.** Components for automated solid-phase synthesis.

| Item | Contents | Dose per well per cycle |
|---|---|---|
| Amino acid plate (96-DWP, 2 mL) | Fmoc-amino acid solutions, 0.21 M in DMF | 100 µL |
| 200 mL PTFE reservoir | HATU, 0.19 M in DMF | 100 µL |
| 200 mL PTFE reservoir | DIPEA, 0.42 M | 100 µL |
| 200 mL PTFE reservoir | Deprotection: 20 % 4-methylpiperidine in DMF | 180 µL × 3 |
| 200 mL PTFE reservoir | DMF for washing | 900 µL × 3 and × 2 |

#### Synthesis cycle and duration

A deck with the appropriate labware was created in the EVOware control software, and a synthesis script was written (S2 Fig). The operator manually enters the synthesis number, number of cycles, incubation times for deprotection, coupling, mixing, and liquid removal, and the path to the directory containing the worklist files that specify the reagent aspiration and dispensing coordinates. The steps from deprotection to the final wash after coupling are looped for the number of couplings specified at the start of the script; after all cycles are complete, a final Fmoc deprotection of the N-terminal residue is performed. The single-residue coupling cycle and its timing breakdown are shown in Fig 7 and Table 2.

**Fig 7.**
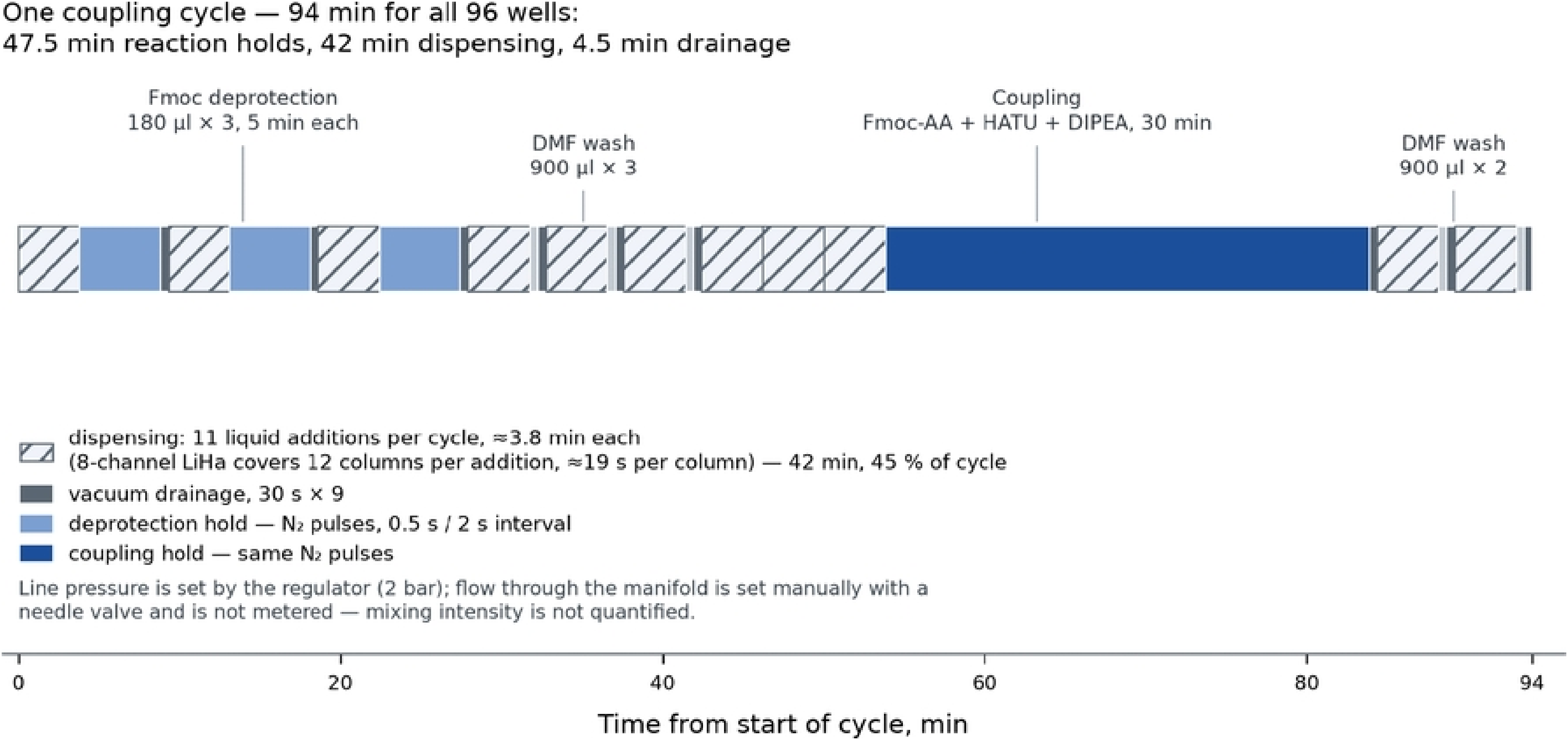
Single-residue coupling cycle and its duration. 94 min per cycle, with 47.5 min of incubation.

**Table 2.** Duration of automated peptide synthesis steps.

| **Element** | **Number per cycle** | **Min per cycle** | **Fraction of cycle** |
|---|---|---|---|
| Robot dispensing (11 dispensing operations × 12 columns) | 11 | 42.0 | 45 % |
| Coupling incubation | 1 | 30.0 | 32 % |
| Deprotection incubation | 3 | 15.0 | 16 % |
| Vacuum drainage (30 s) | 9 | 4.5 | 5 % |
| Wash incubation | 5 | 2.5 | 3 % |
| Coupling cycle |  | 94 | 100 % |

#### Software layer: protocol generation from a request file

Aspiration and dispensing coordinates for every step of each cycle iteration are generated by a separate software component, worklists_creator, which converts a request file into a set of GWL-format worklist files [32–36]. The input is an xlsx table containing 96 sequences and requester names (S2 file PARSYNT_P_inputfile_TEC033.xlsx); the output is a directory of GWL files organized by cycle iteration (step_1.gwl … step_6.gwl for each cycle, and an optional step_cap.gwl) (S3 file PARSYNT_P_worklists_TEC033.zip).

#### Processing logic

For each specified sequence, the C-terminal amino acid residue is excluded because it is already immobilized (preloaded) on the solid support, and the remaining polypeptide chain is reversed in the C → N direction to match the order of synthetic elongation.

The request table is sorted by length and reverse sequence in descending order and distributed across the plate; each well is assigned a barcode of the form TEC + synthesis number + well coordinate (for example, TEC033A01), which then accompanies the sample through analysis.

The request is expanded into a ‘well × amino acid × coupling number’ table, from which a summary of amino acid frequencies is used to calculate solution volumes, numbers of portions, and masses of leophilizated Fmoc derivatives to be weighed out (volume × concentration × molar mass).

Step parameters (labware name in EVOware terminology, volume, solution concentration, and number of repeats) are not hard-coded but declared in the method_proposal.json configuration file for steps 1–6. The protocol is thus described declaratively and can be adjusted without modifying the program.

GWL files are generated using the robotools library [37], with explicit labware definitions (an 8 × 12 reagent plate with a working range of 60–2000 μL and 8 × 1 reagent reservoirs with a range of 60–24 000 μL), in disposable-tip mode. Each file contains one or more dispensing operations per well (96 Aspirate/Dispense commands) and tip-disposal commands. If the required dispensing volume exceeds the maximum permitted tip volume, it is split into portions of the maximum permitted volume, increasing the number of reagent dispensing operations accordingly; ‘С’ comment lines record the source and destination, making the worklist readable and verifiable before execution. The reagent names in these comments are inherited from the configuration and do not affect execution, since the instrument reads only coordinates and volumes. For reagents, one tip serves a row of 12 wells, requiring 8 tips per dispensing operation. Amino acids are dispensed according to a reagent-oriented principle rather than plate topology: a single tip sequentially dispenses a given amino acid into all target wells of the current elongation cycle and is discarded only when the monomer changes. Tip consumption at this stage equals the number of distinct amino acids in the cycle rather than the number of occupied wells: in the first cycle of synthesis TEC033, this is 7 tips for 96 wells. By comparison, dispensing ‘by rows’ would require either 96 tips or washing between wells, while reusing one tip for different amino acids would cause cross-contamination.

Tip consumption is tracked against rack capacity, with five racks of 200 μL tips and one rack of 1000 μL tips per run; when supplies are exhausted, the generator inserts a stop event with an operator message into the protocol, and all such events are compiled in an event log to schedule instrument servicing during synthesis.

Auxiliary files (S4 file PARSYNT_P_auxiliary files_TEC033.zip) include the number of couplings of each amino acid in each sequence synthesized in the batch and the total number of couplings (aa_stat), a solution preparation table calculated to account for dead (non-pipettable) volume and liquid losses during pipetting (aa_suspensions), a log of manual intervention events (events), and a set of synthesis monitoring plots:

● Amino acid consumption: required versus loaded volume for each of the 16 amino acids in synthesis TEC033 (S3 Fig (A));
● Resin layout: the C-terminal residue in each well before synthesis starts (S3 Fig (B));
● Sequence growth: the state of the peptide plate after each cycle, from a single letter to the full sequence (S3 Fig (C));
● Amino acid plate status: the volume remaining in each well after each cycle (initial fill 1800 μL, working positions in columns 01–07); the map shows which positions are approaching depletion and when refilling will be required (S3 Fig (D));
● Manual operation log by cycle: reagent refilling is required in every cycle, and replacement of the 1000 μL tip rack is required in cycles 3 and 5 of seven (S3 Fig (E)).

The generated outputs are not a retrospective report of protocol execution but tools for advance process planning: they contain calculated masses of amino acid derivatives to be weighed out, their placement in the plate wells, and a list of synthesis cycles with scheduled pauses for operator intervention.

The scope of automation covers experimental planning, leaving direct manipulator control to the robot’s native software. The developed algorithm converts request parameters into stoichiometric volumes, component masses to be weighed out, a spatial well layout, and a formalized command sequence. With 96 different sequences and dozens of cycles, this is the main source of manual errors, and without such a software layer the plate format is not viable.

### Protocol verification on the PurePep Chorus synthesizer

#### Model peptide set

A model panel of eight tryptic octapeptides with a C-terminal lysine residue was assembled to validate the protocol. These sequences are proteotypic peptides of blood plasma proteins: terminal components and regulators of the complement system, factors of hemostasis and the contact system, and an extracellular matrix protein (S3 Table).

#### Testing the synthesis protocol

To verify the protocol intended for use on the Tecan liquid handling station, a test synthesis was performed on the PurePep. However, because the minimum dispensing volume of the Chorus is 250 μL, the quantities of all components used were increased 2.5-fold: 25 mg of resin per reactor instead of 10 mg per well, 250 μL of reagent solutions instead of 100 μL, and 2250 μL of washing solvent instead of 900 μL. The molar ratios and excesses relative to the resin were retained.

All eight peptides were obtained and identified. Crude purity was determined as the fraction of the target peak area in the total chromatogram area at 220 nm (S5_File.zip); identity was confirmed by the monoisotopic mass and observation of both charge states, [M+H]⁺ and [M+2H]²⁺ (Table 3).

**Table 3.** Identification and purity of the eight model peptides (PurePep Chorus).

| **Preparation** | **Sequence** | **RT, min** | **Crude purity, %** | **Monoisotopic mass, Da** |
|---|---|---|---|---|
| NP0580101 | NDLISATK | 10.94 | 55.0 | 860.46 |
| NP0580102 | LELLSLSK | 15.43 | 61.4 | 901.55 |
| NP0580201 | VSMPYASK | 8.59 | 81.5 | 881.43 |
| NP0580202 | AVRPGYPK | 7.76 | 89.1 | 886.50 |
| NP0580203 | FTPTETNK | 7.83 | 82.7 | 936.46 |
| NP0580204 | DTPFSQIK | 9.28 | 82.7 | 934.48 |
| NP0580301 | SGFSFGFK | 10.15 | 83.9 | 875.42 |
| NP0580302 | GSSGLEEK | 5.74 | 61.7 | 805.38 |

The selected chemical conditions (HATU/DIPEA, deprotection with 4-methylpiperidine, and a coupling time of 30 min) were found to provide consistent yield and purity of the target peptides in an automated format.

### Test synthesis in a plate (TEC033)

Initial protocol verification on the Chorus platform confirmed successful production of all eight model peptides. In the next stage, this panel was synthesized in parallel in a full-plate format; the results are presented in Fig 8.

**Fig 8.**
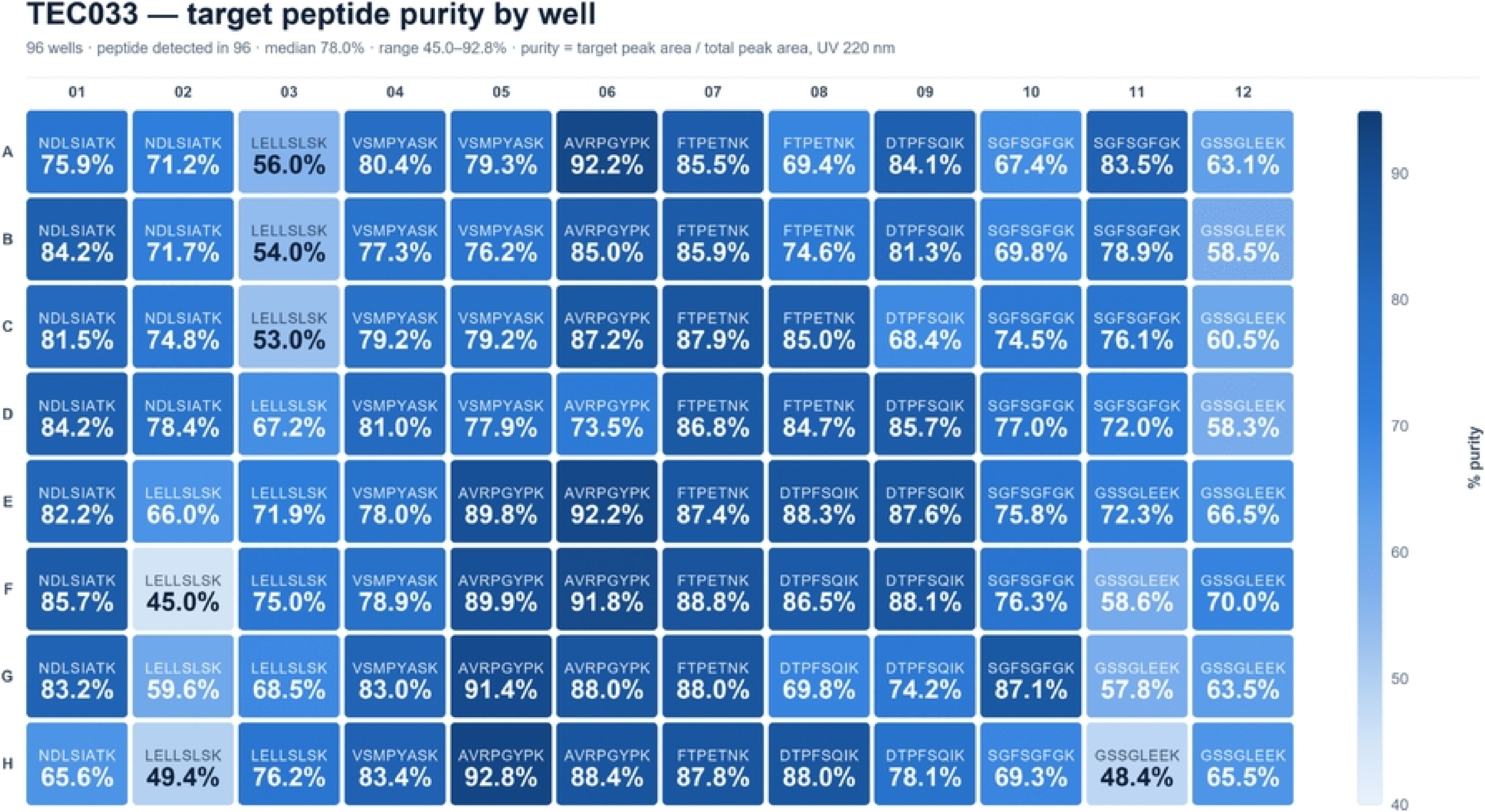
Plate TEC033: crude purity by well.

HPLC–UV–MS analysis (S6_File.zip) showed that the target peptide was detected in all 96 wells; the plate median was 78.0 %, with a range of 45.0 to 92.8 %.

Comparison of the syntheses showed comparable percentages of target peptides in the crude products (Table 4).

**Table 4.** Comparison of crude purity: PurePep Chorus vs plate (TEC033).

| **Peptide** | **Chorus, %** | **Plate, median %** |
|---|---|---|
| NDLISATK | 55.0 | 80.0 |
| LELLSLSK | 61.4 | 62.8 |
| GSSGLEEK | 61.7 | 61.8 |
| VSMPYASK | 81.5 | 79.2 |
| DTPFSQIK | 82.7 | 84.9 |
| FTPTETNK | 82.7 | 86.3 |
| SGFSFGFK | 83.9 | 75.9 |
| AVRPGYPK | 89.1 | 89.8 |

### Uniformity of synthesis across the plate (TEC034)

To assess synthesis uniformity (one peptide × 96 wells), the sequence AVRPGYPK was selected, as it had shown the highest median purity, 89.8 %, with a range of 73.5–92.8 %. In synthesis TEC034, the target peptide was detected in all 96 wells (Fig 9; S7_File.zip).

**Fig 9.**
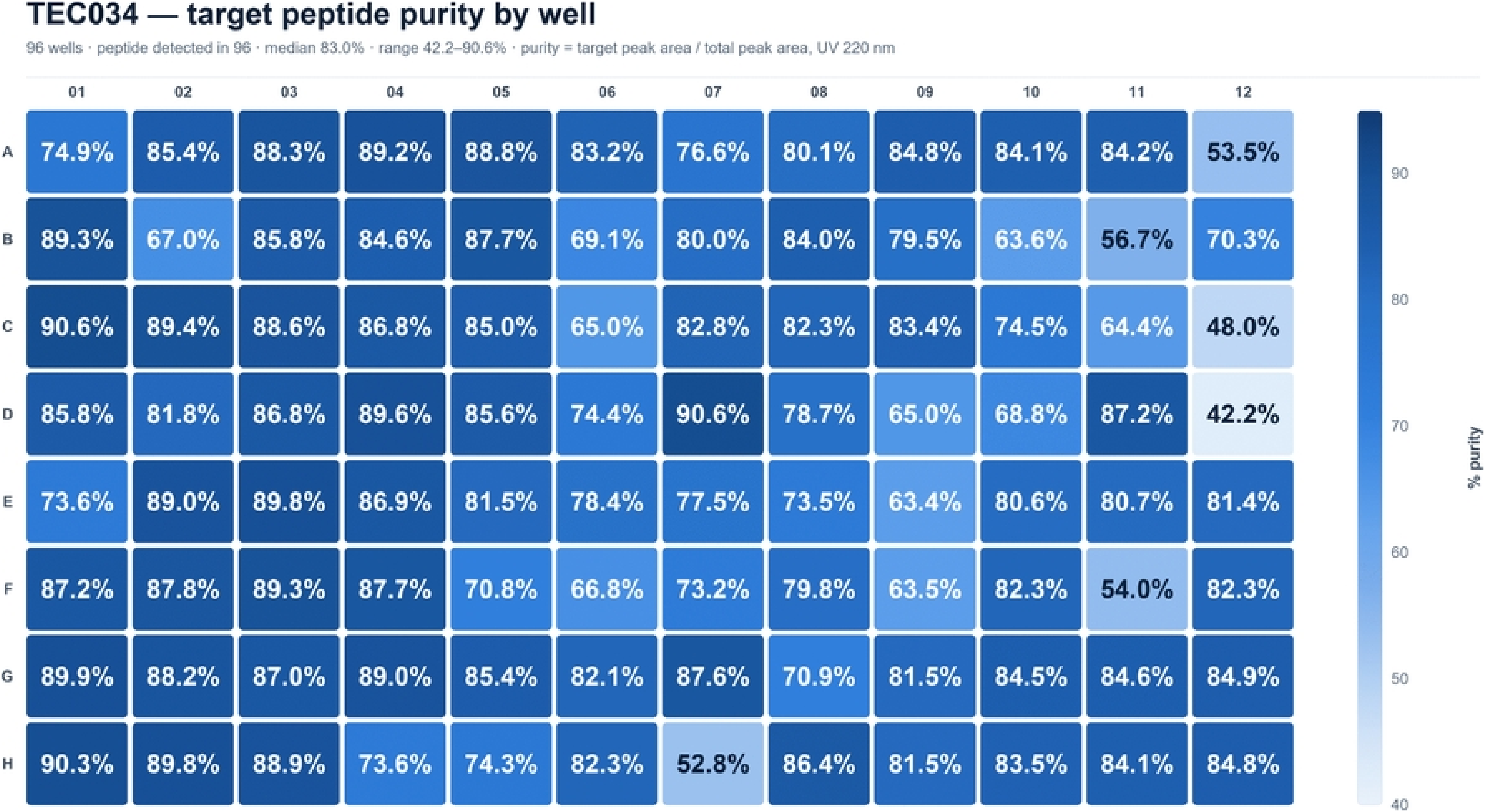
Map of the percentage content of crude AVRPGYPK peptide in each of the 96 wells.

### Preparation of NAT/SIL standards for AVRPGYPK

Because purity assessed by relative peak area does not reflect the absolute amount of substance, and two wells with the same purity may contain different amounts of product, equivalence of positions in terms of absolute content can be established only using a quantitative map based on direct concentration measurements. Quantitative analysis requires a pair of standards of the same sequence: a natural form (NAT) and an isotope-labeled form (SIL, stable isotope-labeled standard), purified and characterized for concentration.

The natural form of AVRPGYPK was synthesized on resin preloaded with C-terminal lysine, and the labeled form on Fmoc-Lys(Boc)-[¹³C₆,¹⁵N₂]-TCP resin. Both forms were produced in preparative quantities and purified. Concentrations were determined by amino acid analysis with MS/MS detection (Table 5). Calibration curves for the amino acids constituting AVRPGYPK are presented in S4 Fig.

**Table 5.**
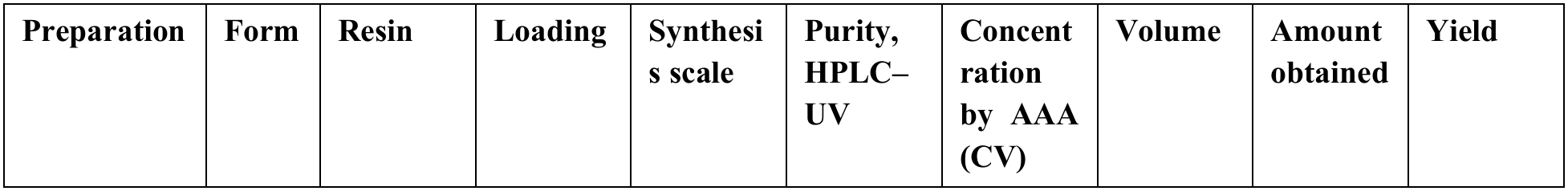
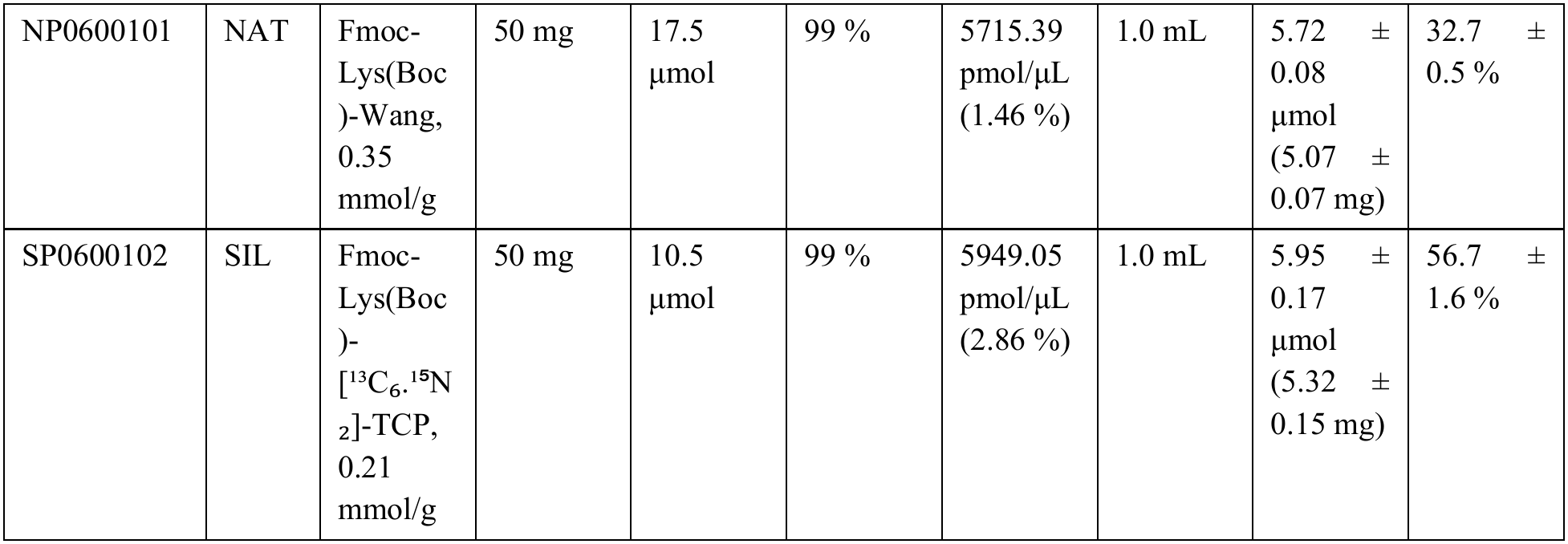
Synthesis, purification, and characterization of AVRPGYPK NAT/SIL standards.

| **Preparation** | **Form** | **Resin** | **Loading** | **Synthesis scale** | **Purity, HPLC–UV** | **Concentration by AAA (CV)** | **Volume** | **Amount obtained** | **Yield** |
|---|---|---|---|---|---|---|---|---|---|
| NP0600101 | NAT | Fmoc-Lys(Boc)-Wang, 0.35 mmol/g | 50 mg | 17.5 µmol | 99 % | 5715.39 pmol/µL (1.46 %) | 1.0 mL | 5.72 ± 0.08 µmol (5.07 ± 0.07 mg) | 32.7 ± 0.5 % |
| SP0600102 | SIL | Fmoc-Lys(Boc)-[<sup>13</sup>C<sub>6</sub>.<sup>15</sup>N<sub>2</sub>]-TCP, 0.21 mmol/g | 50 mg | 10.5 µmol | 99 % | 5949.05 pmol/µL (2.86 %) | 1.0 mL | 5.95 ± 0.17 µmol (5.32 ± 0.15 mg) | 56.7 ± 1.6 % |

### Quantitative plate map

The resulting standards were used to construct calibration curves over the range 2.5–500 pmol/μL (S5 Fig) and to measure AVRPGYPK concentrations in all 96 wells (Fig 10., Table 6, S4). These values were adjusted to the original sample to account for dilution.

**Fig 10.**
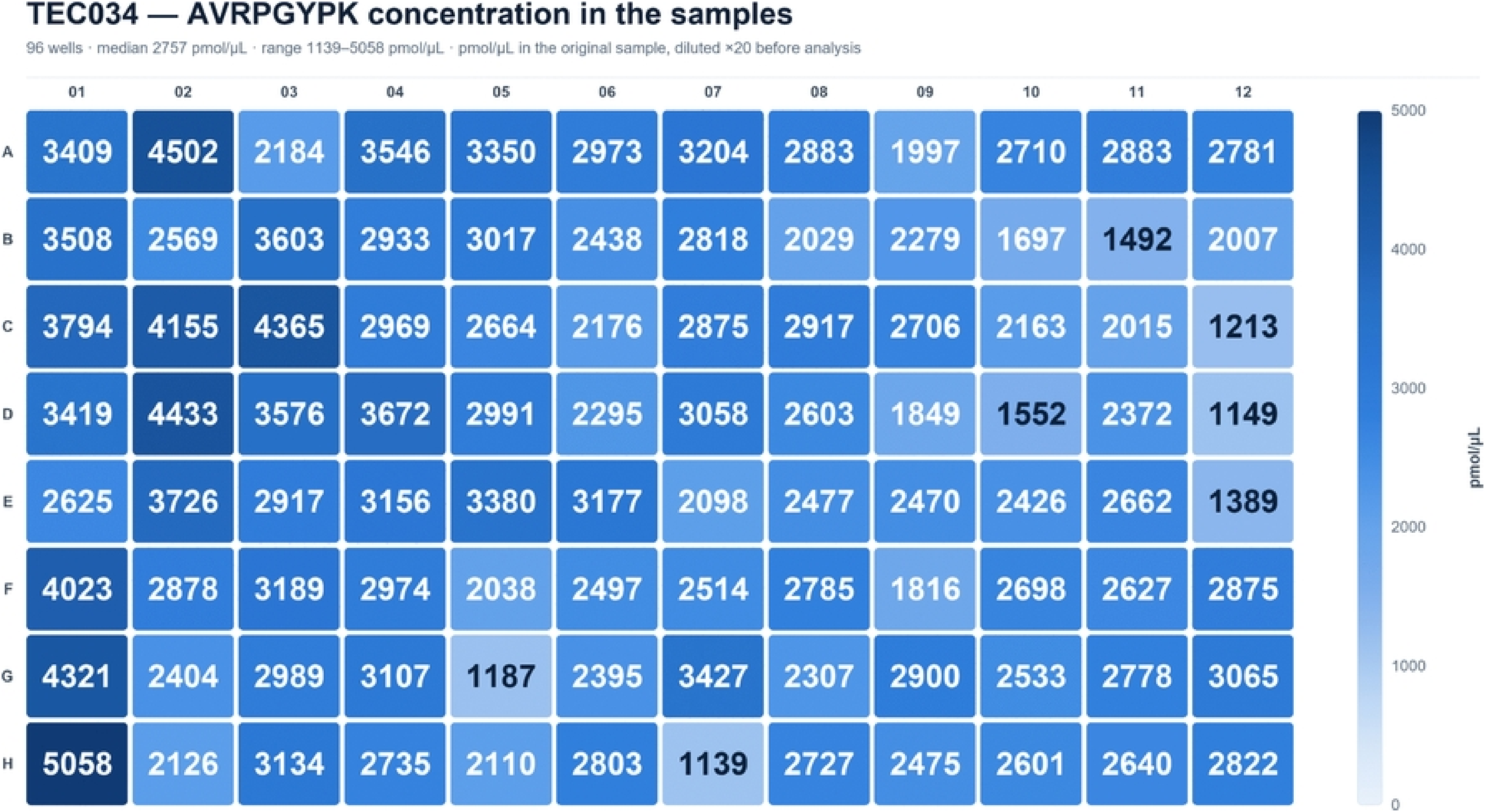
AVRPGYPK concentration by well.

**Table 6.** Target peptide yield per well, TEC034.

| **Parameter (per well)** | **Value** |
|---|---|
| Target peptide | Median 2.76 µmol (2.44 mg) |
| Full range | 1.01–4.48 mg |
| Yield relative to the nominal 3.5 µmol | Median 79 % |
| Total from the plate | 235 mg of target peptide |

The results indicate high synthesis efficiency (median yield 79%, 2.44 mg per well), meaning that peptides obtained in this way can be used not only for candidate selection but also for subsequent certification as standards after purification and amino acid analysis.

### Synthesis of 96 different sequences

The final verification of system operation was the synthesis of 96 different sequences. This simultaneously tested the stated throughput and imposed the most demanding test on the software layer.

The set comprised tryptic peptides 6–9 residues long (S5 Table). In synthesis TEC035, the target peptides were detected in all 96 wells (Fig 11; S8_File.zip).

**Fig 11.**
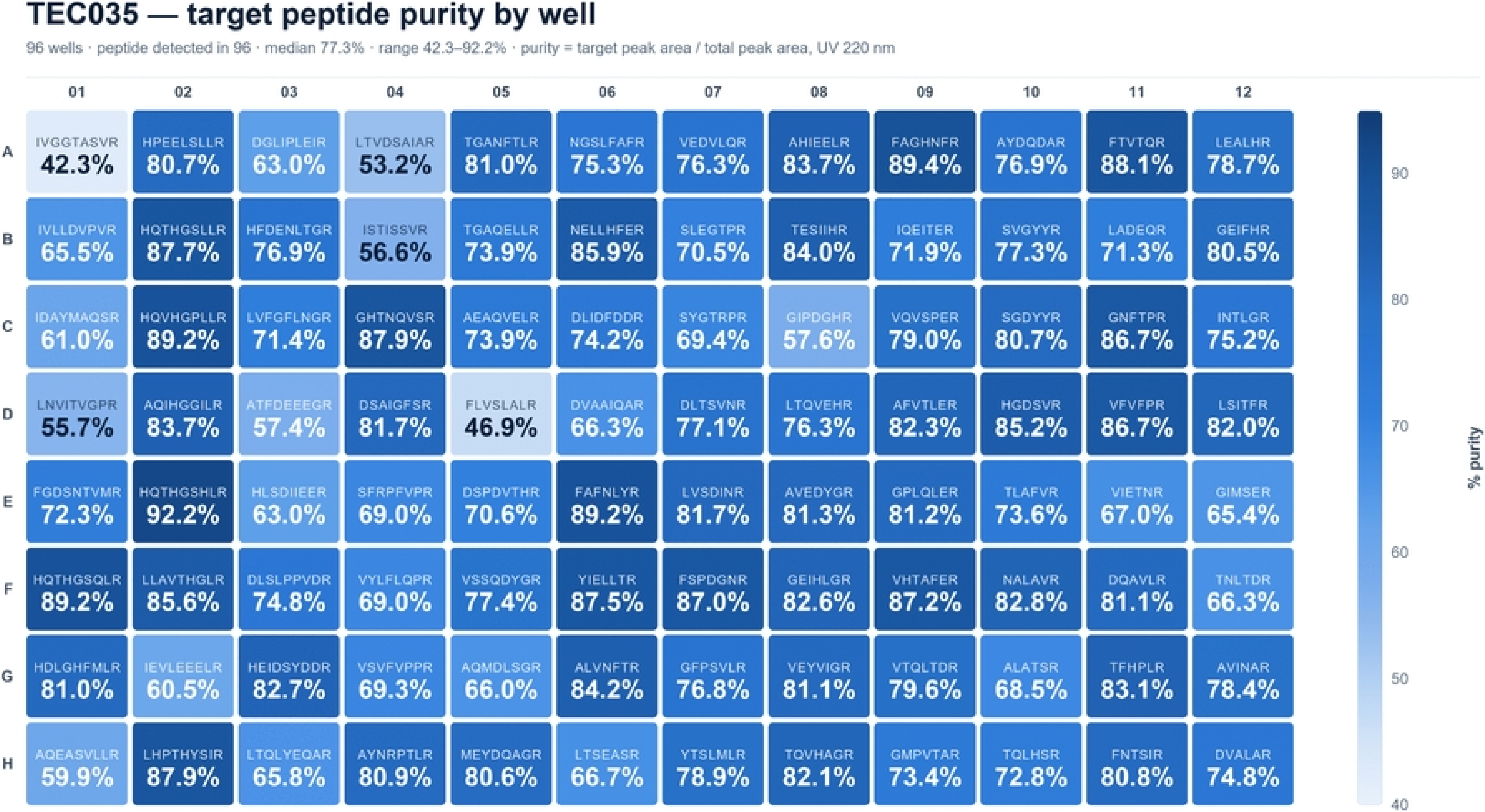
Final plate: 96 different sequences. Median 77.3 %, range 42.3–92.2 %.

The result shows that the stated throughput was achieved in full. The absence of unsynthesized peptides is an important indicator of stable system operation, and the software layer withstood operation at full capacity. Ninety-six syntheses were calculated, executed, and completed without manual intervention.

### Economics: add-on cost and synthesis cost

The add-on bill of materials (S6 Table) contains 13 items; the total cost of purchased components was $351. This sum excludes machining of the manifold and reservoirs with lids, printing of the holder and control unit housing (performed by the coauthors), and the cylinder regulator (already available); the collection vessel was an empty HPLC solvent bottle.

The synthesis cost calculation comprises two main items: tips and chemical reagents. Labor costs for staff involved in peptide synthesis were not included.

Tip consumption is calculated directly from the generated worklist files (Table 7), since it depends solely on the dispensing scheme. Within a chain elongation cycle (11 operations per plate), common solutions (HATU, DIPEA, 20% 4-methylpiperidine, and DMF) are dispensed using 8 tips per operation (one tip per row of 12 wells), whereas amino acid tip consumption is determined by the number of unique monomers in the current cycle through grouped dispensing.

**Table 7.** Tip consumption per coupling cycle.

| **Dispensing** | **Volume** | **Repeats per cycle** | **Tip type** | **Tips per cycle** |
|---|---|---|---|---|
| Deprotection (step_1) | 180 µL | 3 | 200 µL | 24 |
| DMF wash (step_2) | 900 µL | 3 | 1000 µL | 24 |
| Amino acid (step_3) | 100 µL | 1 | 200 µL | Equal to the number of distinct amino acids |
| HATU (step_4) | 100 µL | 1 | 200 µL | 8 |
| DIPEA (step_5) | 100 µL | 1 | 200 µL | 8 |
| DMF wash (step_6) | 900 µL | 2 | 1000 µL | 16 |

With a plate fully loaded with different sequences (up to 20 distinct amino acids per cycle), consumption is 60 tips of 200 μL and 40 tips of 1000 μL per cycle, costing $11.53 per plate, or $0.120 per coupling in a well (S7 Table).

It should be noted that tip costs can be avoided if the liquid-handler station is equipped with reusable tips that are washed after each dispensing step.

The second cost item is chemical reagents (S8 Table), which are consumed in each cycle, so their cost is proportional to the number of couplings. Resin, cleavage cocktail, and final N-terminal Fmoc deprotection (including tips for this step) are used only once, and their cost does not depend on peptide length. This gives the following formula for coupling cost per well: (reagents $0.109 + tips $0.120) + (final Fmoc deprotection ($0.054 reagents + $0.058 tips) / n-cycles) = $0.229 + ($0.1128 / n-cycles). The cost of a peptide is then calculated as: resin + cleavage + (n-cycles × $0.229 + final Fmoc deprotection ($0.054 reagents + $0.058 tips), and should be reported for specific lengths. Fig 12a demonstrates the dependence of peptide cost on its length. The increase in cost is due solely to increased consumption of consumables (reagents + tips).

**Fig 12.**
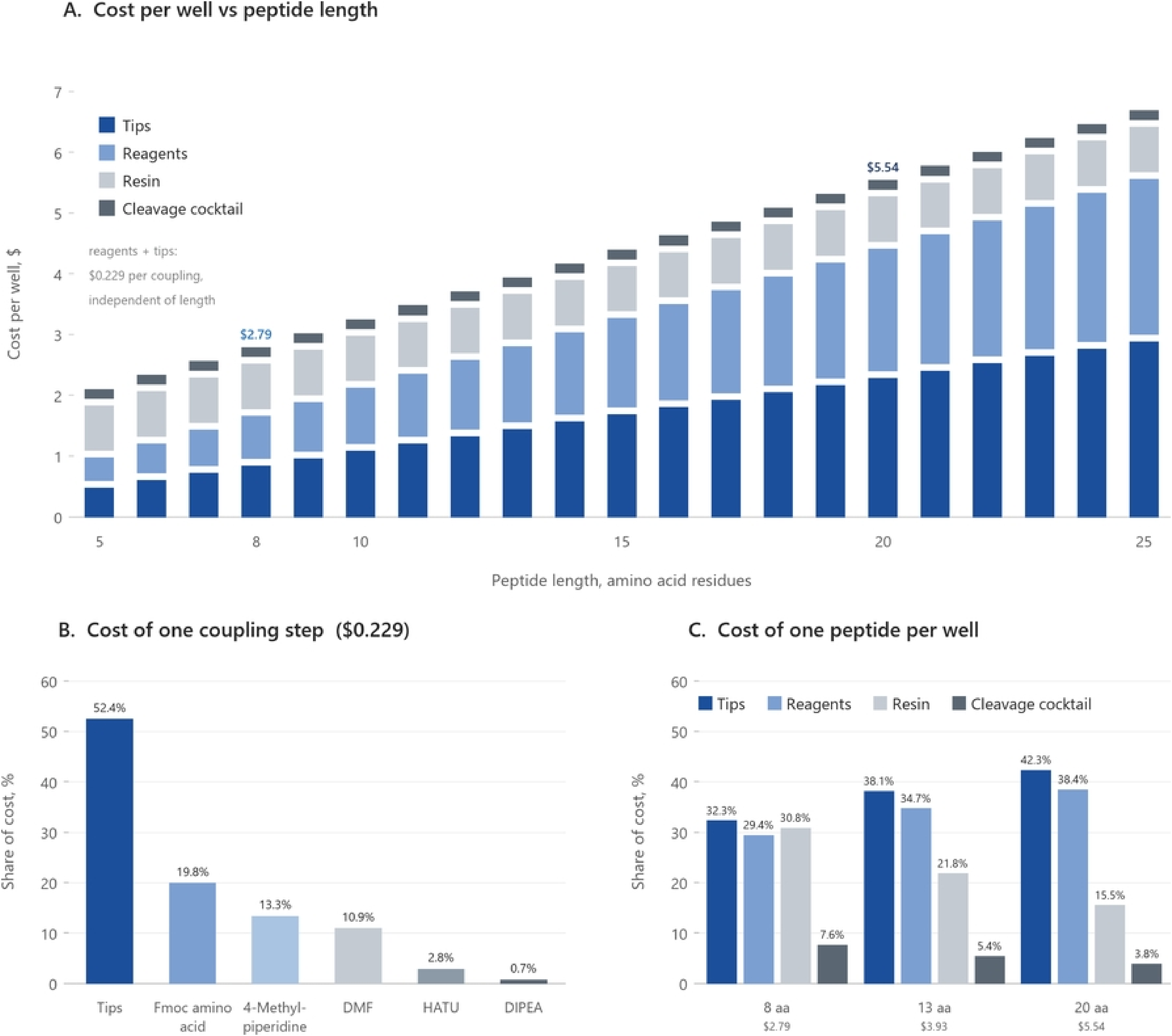
Cost of peptide synthesis. (A) Cost per well versus peptide length, with the segments representing tips, reagents, resin and cleavage cocktail. Per-step reagents and tips grow linearly with the number of coupling steps ($0.229 per coupling step); the final Fmoc deprotection, resin (10 mg per well) and cleavage cocktail (1 mL per well) are one-off costs, so their share falls with peptide length. (B) Percentage composition of a single coupling step (excluding one-off costs). (C) Percentage composition of the total cost of one peptide per well for 8, 13 and 20 residues.

### Comparison with open and commercial solutions

Analysis and comparison of similar systems (S9 Table) showed that each was developed for a specific task and addresses only that task: for example, synthesizer by Kallmyer et al. (Frugal) [22] is an extremely inexpensive automated peptide synthesizer for individual syntheses, PepSy [21] is primarily an affordable stand-alone synthesizer, Chemputer [16] formalizes complex chemistry and completely removes manual work from the pipeline, TaPSy [6] enables parallel production of 12 different sequences on a scale of hundreds of milligrams, and the 384-well SPPS system reported by Schüttel et al. [20] enables the creation of extensive libraries (1536 at a time) of short peptides (2-3 amino acid residues). ParSynt-P solves a task that none of these systems addresses: obtaining many different sequences of 6 or more amino acid residues in microgram-to-milligram quantities, inexpensively and rapidly, as needed to select candidates for peptide standards. A $351 add-on for an existing robot provides 96 positions per run at $0.23 per coupling, so synthesis cost ceases to be a factor limiting the size of the candidate panel. The boundaries of these niches are shown in Fig 13.

**Fig 13.**
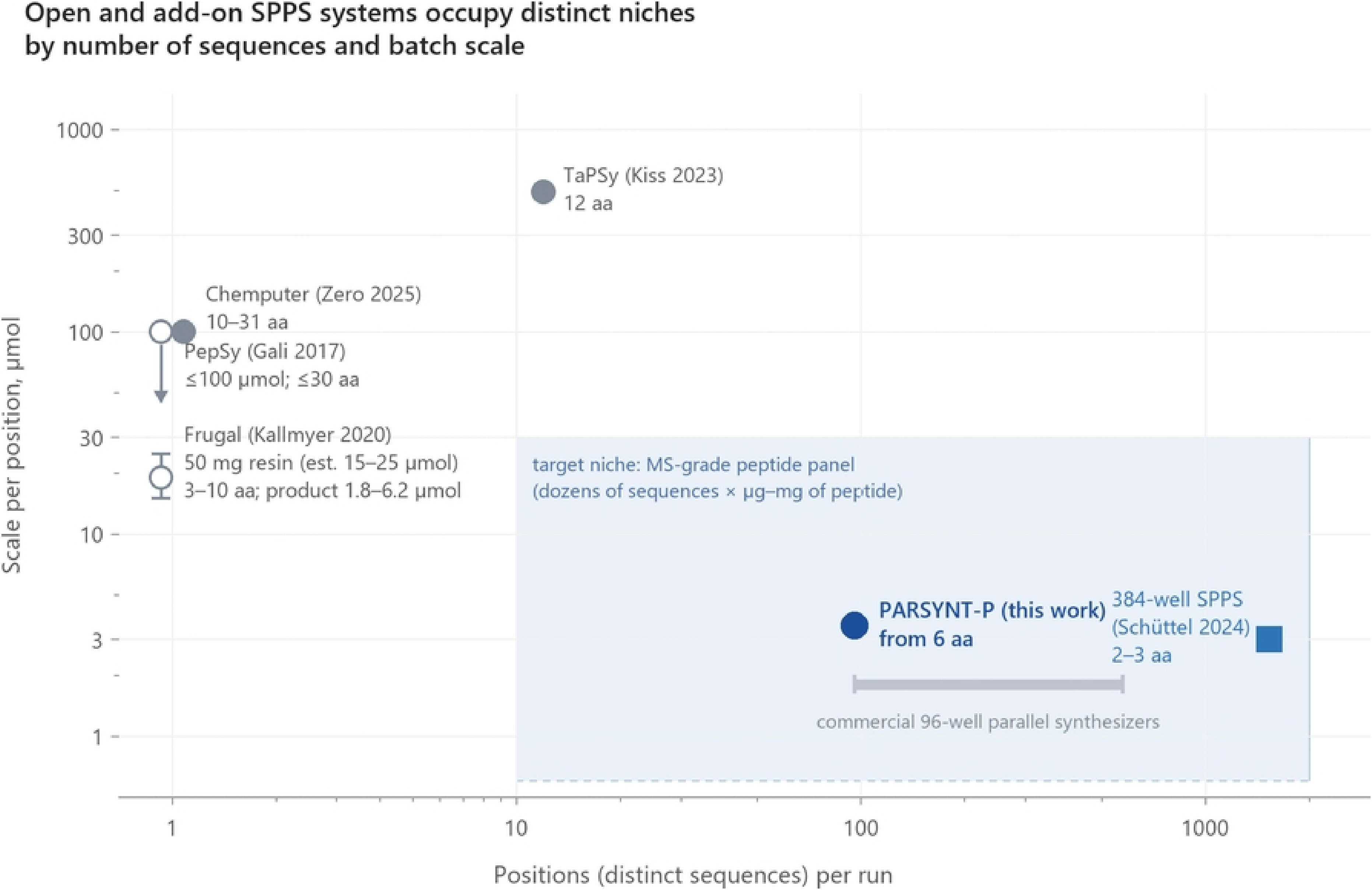
Niche map: open and add-on SPPS systems. Distribution of systems by number of parallel sequences, loading scale, and sequence length.

## Conclusions

This work demonstrates that the developed hardware and software system enables a standard liquid handling station in a proteomics laboratory to be upgraded to a high-throughput parallel plate-format peptide synthesizer, eliminating the need to purchase specialized commercial equipment. System operation was confirmed on three full-format 96-well plates. The target product was successfully obtained in all 288 wells, a key reliability criterion for generating panels of standards. In a comparative analysis using an identical set of eight sequences, the developed plate format was not inferior to the commercial reference synthesizer. Quantitative characterization made it possible to determine the median yield across the plate, which was 2.44 mg of target peptide per well. The synthesis cost ($0.23 per coupling and $1.14 per milligram of crude product) is an order of magnitude lower than commercial alternatives, while the cost of hardware components is comparable to that of the lowest-cost open platforms at a synthesis capacity two orders of magnitude greater.

## Acknowledgments

This manuscript was edited for English language and stylistic flow using an AI tool. Some of the figures and all of the chromatograms were compiled by an AI tool based on raw data.

## Supporting information

**S1 Fig. Manifold and stand assembly drawing (two sheets).**

**S2 Fig. Synthesis script in EVOware.**

**S3 Fig. Synthesis monitoring plots (a–e).**

**S4 Fig. Amino acid calibration curves for AVRPGYPK.**

**S5 Fig. LC–MS/MS calibration curves for AVRPGYPK.**

**S1 Table. MRM transitions for amino acid analysis.**

**S2 Table. MRM transitions for quantification of AVRPGYPK.**

**S3 Table. Model peptide set synthesized on the PurePep Chorus.**

**S4 Table. TEC034: purity, concentration, and yield across 96 wells.**

**S5 Table. TEC035: plate composition and the 96 sequences.**

**S6 Table. Bill of materials.**

**S7 Table. Tip consumption and cost.**

**S8 Table. Reagent prices and cost per synthesis cycle (sheets S8a–S8d).**

**S9 Table. Comparison of automated SPPS systems. ParSynt-P source code.**

**S1 File Cotroller_mod.zip**

**S2 file PARSYNT_P_inputfile_TEC033.xlsx**

**S3 file PARSYNT_P_worklists_TEC033.zip**

**S4 file PARSYNT_P_auxiliary files_TEC033.zip**

**S5 File.zip**

**S6 File.zip**

**S7 File.zip**

**S8 File.zip**

## Notes

### Competing Interest Statement

The authors have declared no competing interest.

## References

1. Picotti P, Rinner O, Stallmach R, Dautel F, Farrah T, Domon B, et al. High-throughput generation of selected reaction-monitoring assays for proteins and proteomes. Nature Methods. 2009;7(1):43–46. doi:10.1038/nmeth.1408

2. Zolg DP, Wilhelm M, Schnatbaum K, Zerweck J, Knaute T, Delanghe B, et al. Building ProteomeTools based on a complete synthetic human proteome. Nature Methods. 2017;14(3):259–262. doi:10.1038/nmeth.4153

3. Vakaryuk AV, Butenko IO, Kitsilovskaya NA, Kovalenko AV, Gremyacheva VD, Kozhemyakin GL, et al. Implementation of an Internalized Hemolysis Assessment Approach in Targeted Bottom-Up Proteomics for the Purpose of Blood Plasma Sample Quality Control. Journal of Proteome Research. 2026. doi:10.1021/acs.jproteome.6c00157

4. Kuzyk MA, Smith D, Yang J, Cross TJ, Jackson AM, Hardie DB, et al. Multiple Reaction Monitoring-based, Multiplexed, Absolute Quantitation of 45 Proteins in Human Plasma. Molecular & Cellular Proteomics. 2009;8(8):1860–1877. doi:10.1074/mcp.m800540-mcp200

5. Percy AJ, Borchers CH. Detailed Method for Performing the ExSTA Approach in Quantitative Bottom-Up Plasma Proteomics. Methods in Molecular Biology. 2021:353–384. doi:10.1007/978-1-0716-1024-4_25

6. Kiss K, Ránky S, Gyulai Z, Molnár L. Development of a novel, automated, robotic system for rapid, high-throughput, parallel, solid-phase peptide synthesis. SLAS Technology. 2023;28(2):89–97. doi:10.1016/j.slast.2023.01.002

7. Mäde V, Els-Heindl S, Beck-Sickinger AG. Automated solid-phase peptide synthesis to obtain therapeutic peptides. Beilstein Journal of Organic Chemistry. 2014;10:1197–1212. doi:10.3762/bjoc.10.118

8. Merrifield RB. Solid Phase Peptide Synthesis. I. The Synthesis of a Tetrapeptide. Journal of the American Chemical Society. 1963;85(14):2149–2154. doi:10.1021/ja00897a025

9. Amblard M, Fehrentz JA, Martinez J, Subra G. Methods and Protocols of Modern Solid Phase Peptide Synthesis. Molecular Biotechnology. 2006;33(3):239–254. doi:10.1385/mb:33:3:239

10. Behrendt R, White P, Offer J. Advances in Fmoc solid-phase peptide synthesis. Journal of Peptide Science. 2016;22(1):4–27. doi:10.1002/psc.2836

11. Simon MD, Heider PL, Adamo A, Vinogradov AA, Mong SK, Li X, et al. Rapid Flow-Based Peptide Synthesis. ChemBioChem. 2014;15(5):713–720. doi:10.1002/cbic.201300796

12. Mijalis AJ, Thomas DA, Simon MD, Adamo A, Beaumont R, Jensen KF, et al. A fully automated flow-based approach for accelerated peptide synthesis. Nature Chemical Biology. 2017;13(5):464–466. doi:10.1038/nchembio.2318

13. Hartrampf N, Saebi A, Poskus M, Gates ZP, Callahan AJ, Cowfer AE, et al. Synthesis of proteins by automated flow chemistry. Science. 2020;368(6494):980–987. doi:10.1126/science.abb2491

14. Collins JM, Singh SK, White TA, Cesta DJ, Simpson CL, Tubb LJ, et al. Total wash elimination for solid phase peptide synthesis. Nature Communications. 2023;14(1). doi:10.1038/s41467-023-44074-5

15. Pacini L, Muthyala M, Aguiar L, Zitterbart R, Rovero P, Papini AM. Optimization of peptide synthesis time and sustainability using novel eco-friendly binary solvent systems with induction heating on an automated peptide synthesizer. Journal of Peptide Science. 2024;30(9). doi:10.1002/psc.3605

16. Zero J, Tyler TJ, Cronin L. Universal peptide synthesis via solid-phase methods fused with chemputation. Nature Communications. 2025;16(1). doi:10.1038/s41467-025-62344-2

17. Frank R. Spot-synthesis: an easy technique for the positionally addressable, parallel chemical synthesis on a membrane support. Tetrahedron. 1992;48(42):9217–9232. doi:10.1016/s0040-4020(01)85612-x

18. Loeffler FF, Foertsch TC, Popov R, Mattes DS, Schlageter M, Sedlmayr M, et al. High-flexibility combinatorial peptide synthesis with laser-based transfer of monomers in solid matrix material. Nature Communications. 2016;7(1). doi:10.1038/ncomms11844

19. Mehta AY, Veeraiah RKH, Dutta S, Goth CK, Hanes MS, Gao C, et al. Parallel Glyco-SPOT Synthesis of Glycopeptide Libraries. Cell Chemical Biology. 2020;27(9):1207–1219.e9. doi:10.1016/j.chembiol.2020.06.007

20. Schüttel M, Will E, Sangouard G, Zarda A, Habeshian S, Nielsen AL, et al. Solid-phase peptide synthesis in 384-well plates. Journal of Peptide Science. 2024;30(4). doi:10.1002/psc.3555

21. Gali H. An Open-Source Automated Peptide Synthesizer Based on Arduino and Python. SLAS Technology. 2017;22(5):493–499. doi:10.1177/2472630316685844

22. Kallmyer NE, Rider NE, Reuel NF. Design and validation of a frugal, automated, solid-phase peptide synthesizer. PLOS ONE. 2020;15(8):e0237473. doi:10.1371/journal.pone.0237473

23. Anderson NL, Anderson NG. The Human Plasma Proteome. Molecular & Cellular Proteomics. 2002;1(11):845–867. doi:10.1074/mcp.r200007-mcp200

24. Geyer PE, Kulak NA, Pichler G, Holdt LM, Teupser D, Mann M. Plasma Proteome Profiling to Assess Human Health and Disease. Cell Systems. 2016;2(3):185–195. doi:10.1016/j.cels.2016.02.015

25. Picotti P, Aebersold R. Selected reaction monitoring–based proteomics: workflows, potential, pitfalls and future directions. Nature Methods. 2012;9(6):555–566. doi:10.1038/nmeth.2015

26. Gillette MA, Carr SA. Quantitative analysis of peptides and proteins in biomedicine by targeted mass spectrometry. Nature Methods. 2013;10(1):28–34. doi:10.1038/nmeth.2309

27. Mallick P, Schirle M, Chen SS, Flory MR, Lee H, Martin D, et al. Computational prediction of proteotypic peptides for quantitative proteomics. Nature Biotechnology. 2006;25(1):125–131. doi:10.1038/nbt1275

28. Kusebauch U, Campbell DS, Deutsch EW, Chu CS, Spicer DA, Brusniak MY, et al. Human SRMAtlas: A Resource of Targeted Assays to Quantify the Complete Human Proteome. Cell. 2016;166(3):766–778. doi:10.1016/j.cell.2016.06.041

29. Gerber SA, Rush J, Stemman O, Kirschner MW, Gygi SP. Absolute quantification of proteins and phosphoproteins from cell lysates by tandem MS. Proceedings of the National Academy of Sciences. 2003;100(12):6940–6945. doi:10.1073/pnas.0832254100

30. Kirkpatrick DS, Gerber SA, Gygi SP. The absolute quantification strategy: a general procedure for the quantification of proteins and post-translational modifications. Methods. 2005;35(3):265–273. doi:10.1016/j.ymeth.2004.08.018

31. Qasrawi DO, Petrotchenko EV, Borchers CH. Amino acid analysis for peptide quantitation using reversed-phase liquid chromatography combined with multiple reaction monitoring mass spectrometry. Analytical and Bioanalytical Chemistry. 2023;415(22):5261–5267. doi:10.1007/s00216-023-04840-2

32. Kluyver T, Ragan-Kelley B, Pérez F, Granger B, Bussonnier M, Frederic J, et al. Jupyter Notebooks – a publishing format for reproducible computational workflows. In: Positioning and Power in Academic Publishing: Players, Agents and Agendas. IOS Press; 2016. p. 87–90. doi:10.3233/978-1-61499-649-1-87

33. Harris CR, Millman KJ, van der Walt SJ, Gommers R, Virtanen P, Cournapeau D, et al. Array programming with NumPy. Nature. 2020;585(7825):357–362. doi:10.1038/s41586-020-2649-2

34. The pandas development team. pandas-dev/pandas: Pandas [software]. Version v3.1.0rc0. Zenodo; 2026. doi:10.5281/zenodo.3509134

35. Hunter JD. Matplotlib: A 2D Graphics Environment. Computing in Science & Engineering. 2007;9(3):90–95. doi:10.1109/MCSE.2007.55

36. Waskom M. seaborn: statistical data visualization. Journal of Open Source Software. 2021;6(60):3021. doi:10.21105/joss.03021

37. Osthege M, Helleckes L, Nießer J, Prigolovkin L, Halle L, Latour T, et al. JuBiotech/robotools: v1.16.0 [software]. Zenodo; 2026. doi:10.5281/zenodo.4697605

